# Monolayer, Spheroid, and Organoid Formats Elicit Proteomic States that Couple Organelle Organization to Metabolic Vulnerabilities

**DOI:** 10.64898/2026.11.20.768195

**Authors:** Krystal K. Lum, Oscar J. Pundel, William A. Hofstadter, John M. Muroski, Ishea A. Johnson, Benjamin G. Neel, Ileana M. Cristea

**Affiliations:** Department of Molecular Biology, Princeton University, Washington Road, Princeton, NJ; Laura and Isaac Perlmutter Cancer Center, New York University Grossman School of Medicine, New York, NY

**Author notes:** Corresponding author: Ileana M. Cristea, 210 Lewis Thomas Laboratory Department of Molecular Biology Princeton University, Princeton, NJ 08544.

**Keywords:** Organoid, spheroid, mass spectrometry, proteomics, high-grade serous tubo ovarian cancer, membrane contact site

## Abstract

Three-dimensional (3D) cell culture systems are becoming more widely applied to model tissue organization, disease progression, and therapeutic response. Yet, the molecular characteristics that distinguish organoids and spheroids from conventional monolayer culture are incompletely defined. Here, using tumorigenic organoid models of high-grade serous tubo-ovarian cancer (HGSC), we establish a quantitative proteomic resource that reveals how cell architecture and media independently but jointly shape the cellular state in organoids, spheroids, and monolayers. We further pursue HGSC subtype- enriched features within this framework and identify that a poor prognosis-associated genotype with reported paclitaxel hypersensitivity, *Trp53^-/-^;Pten^-/-^;Nf1^-/-^*(PPN), is marked by increased microtubule- associated proteins, lipid-associated factors, and mitochondrial remodeling. PPN cells displayed clustered lipid droplets, whereby mitochondria-ER tethering increased. Relative to *Trp53^-/-^* (P) and a frequent HGSC genotype, *Trp53*^-/-^;*Brca1*^+/-^;*Myc*^OE^ (PBM), live-cell imaging showed that PPN exhibits the highest degree of stable mitochondria-ER encapsulations, coincident with fission-biased mitochondria and a high-flux respiratory state. Disruption of the mitochondria-ER contact protein PTPIP51 reduced lipid droplet clustering in the PPN state. Because cytomegalovirus is known to exploit host mitochondria remodeling, mitochondria-ER contacts, and metabolic output during infection, we tested whether this pre-existing host configuration is associated with cytomegalovirus permissiveness. PPN organoids exhibited greater murine cytomegalovirus infection and infectious output than matched monolayers or PBM. Together, these data define a resource of distinct protein characteristics between matched monolayer, spheroid, and organoid systems that should be considered for experimental design. We also uncover a mechanistic axis connecting 3D architecture, organelle contacts, metabolism, drug response, and viral permissiveness.

## Introduction

Three-dimensional (3D) cell and tissue culture models have become increasingly used experimental systems for modeling tissue architecture, disease progression, and therapeutic response^1–3^. Their physiological resemblance to tissues compared with conventional two-dimensional (2D) monolayer cultures is are widely recognized. Organoids exhibit major differences in physical organization and environmental context, as well as more closely approximate multicellular organization, apical-basal polarity, cell-cell and cell-matrix interactions with increased tension, and diffusion-limited gradients of oxygen, nutrients, metabolites, and drugs^4–7^. By contrast, 2D monolayers force cells into a flattened geometry on stiff planar substrates, reduce higher-order matrix engagement, and expose cells relatively uniformly to soluble cues, conditions that can alter cell morphology, polarity, proliferation, and metabolism relative to tissue-like states^2, 4, 8^. Among 3D model systems, suspension spheroids experience cell compaction, altered diffusion, and limited matrix engagement, whereas organoids are typically embedded within extracellular matrix and exhibit matrix-dependent growth, remodeling, and polarity. These differences imply that each 3D growth system models biologically distinct states, yet the molecular features that underlie these differences, and their functional consequences, remain incompletely understood^1–3, 9^.

In many comparative studies of 2D monolayers with either 3D organoids or spheroids, architecture and media are changed simultaneously, confounding whether observed molecular differences arise from physical organization, extracellular signals, or their combinatorial interactions/effects. This issue is relevant in tumor models, where microenvironmental context and tissue geometry can reshape metabolism, organelle organization, and therapeutic responses^8, 10^. Indeed, differences in epithelial trafficking, mitochondrial and metabolic state, and even subcellular ultrastructure have been reported in cells maintained in 2D versus 3D growth conditions^2, 8, 10–12^. A resource that distinguishes architecture-driven from media-driven characteristics would be useful not only for a specific disease context, but also broadly for biology applications that rely on 3D cell models. We also lack a global reference that benchmarks shared versus format-specific states across monolayer, spheroid, and organoid systems within a source-matched and media-controlled experimental framework. Such a resource could be used to interpret why phenotypes differ across experimental formats.

In this study, we leveraged organoid models of high-grade serous tubo-ovarian carcinoma (HGSC). HGSC remains the deadliest ovarian epithelial cancer, as approximately 70-90% of patients die from their disease and survival has been extended modestly over the last several decades^13–15^. Although initial responses to platinum and taxane (e.g., paclitaxel) therapy are common, most advanced tumors recur and ultimately acquire drug-resistant disease, creating a critical need to define subtype-specific vulnerabilities and state-dependent liabilities^12, 13, 16, 17^. We previously generated human genomics-informed syngeneic mouse organoids isolated from Fallopian tube epithelium, the HGSC cell-of-origin^15, 18–21^. The organoids were engineered to represent frequently seen HGSC genotypes, including *Trp53*^-/-^;*Brca1*^+/-^;*Myc*^OE^ (PBM) and *Trp53*^-/-^;*Pten*^-/-^;*Nf1*^-/-^ (PPN)^15, 16, 22–24^. When implanted into syngeneic mice, PPN tumors form with greater penetrance and shorter latency, produce more ascites and omental metastases, and shorten host survival relative to other genotypes, while PTEN and NF1 loss have both been associated with poor outcome in HGSC^15, 16^. PPN also exhibits paclitaxel hypersensitivity relative to PBM and other HGSC subtypes^15^, creating an opportunity to identify liabilities that coexist with aggressive disease behavior. Moreover, cells from these organoids can be adapted to grow as 2D monolayers and as 3D spheroids, providing a tractable system for disentangling the contributions of genotype, architecture, and growth media to cellular state in a genetically defined background.

We exploited this platform to ask how architecture and media jointly determine cell characteristics across monolayers, spheroids, and organoids. We used proteomics to define architecture-specific states, as well as identify and test functional consequences in follow-up experimentation. First, we generated a resource that annotates architecture-dependent and media-controlled proteomic characteristics, identifying format-specific 2D and 3D protein states. Second, we functionally interrogated characteristics elevated in organoids over 2D monolayers. Increased microtubule and microtubule associated protein levels, in conjunction with microscopy analyses of microtubule bundling expand our understanding of features linked to enhanced responsiveness to paclitaxel. We further demonstrate that a protein program involving inter-organelle membrane contact sites, mitochondrial metabolism, and lipid droplet organization is coincident with a particularly poor prognosis subtype. We find that these characteristics are linked with enhanced permissiveness to a metabolically active virus associated with oncomodulation, murine cytomegalovirus (CMV), which is known to remodel host organelles and membrane contact sites, including mitochondrial architecture and mitochondria-ER contacts. In doing so, we provide a general framework for interpreting organoid and spheroid systems relative to conventional 2D monolayers and a mechanistic perspective on how 3D cellular organization shapes function.

## Results

### Ovarian cancer models reveal organoid-specific proteome remodeling relative to 2D monolayers

We first established a genetically defined Fallopian tube epithelial (FTE) platform in which growth architecture, media context, and genotype could be interrogated in a controlled manner (Figure 1A). Starting from parental *Trp53^-/-^* cells and two transformed HGSC-like lines, *Trp53^-/-^;Brca1^+/-^;Myc^OE^* (PBM) and *Trp53^-/-^;Pten^-/-^;Nf1^-/-^* (PPN), we generated matched 2D monolayer and 3D organoid cultures for quantitative proteomics, microscopy, and metabolic assays. This setup provided a resource-level comparison of 2D versus 3D growth and a tractable system for subsequent genotype-dependent mechanistic studies. Confocal imaging confirmed that PBM and PPN cells generate distinct 2D and 3D morphologies, including differences in cytoplasmic surface area and tissue-like organoid volume (Figure 1B). In line with prior findings^15^, these observations support that there are inherent distinct molecular features that distinguish their tumorigenesis and chemosensitivity. As the organoids are embedded in Matrigel®, a basement membrane extract, an important consideration and impediment in interpreting proteomic analyses is the possibility of residual matrix protein carryover from Matrigel® dominating identifications and enriched GO terms. Preliminary experimentation on organoid recovery resulted in a protocol that minimized matrix protein carryover (Figure S1A). Relative abundance distributions showed that the matrix-related proteins occupy a quantitatively similar range between organoids and monolayers. This observation suggests that the organoid state reflects proteome remodeling, as opposed to being driven by matrix carryover.

**Figure 1.**
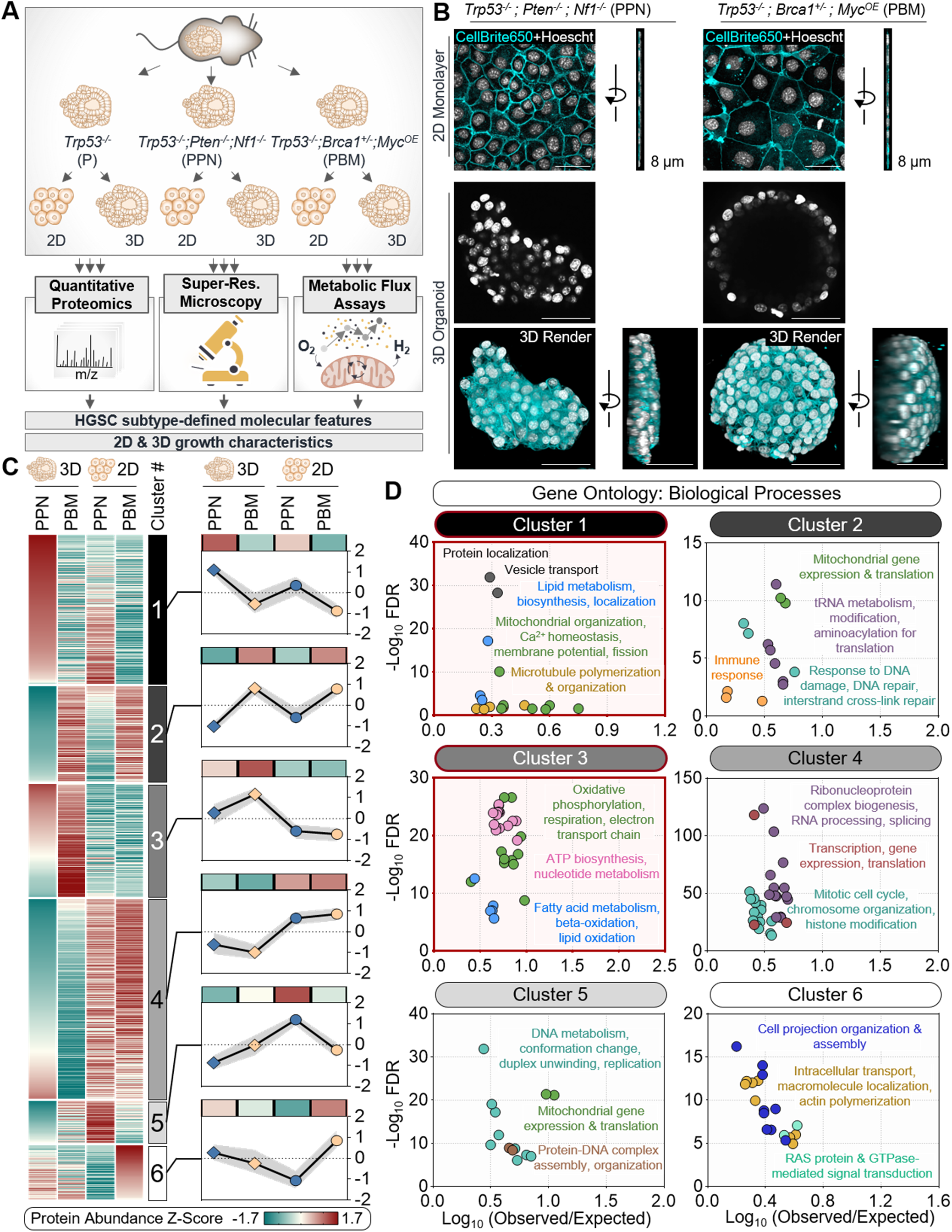
Ovarian cancer models reveal organoid-specific proteome remodeling relative to 2D monolayers. A) Experimental scheme to characterize 2D monolayer and 3D organoid growth characteristics and the molecular features of HGSC subtypes. Parental *Trp53^-/-^*(P), *Trp53^-/-^;Pten^-/-^;Nf1^-/-^*(PPN), and *Trp53^-/-^;Brca1^+/-^;Myc^OE^*(PBM) epithelial models were grown as 2D monolayers and 3D organoids. Samples were subjected to global and targeted quantitative proteomics, and orthogonal assays including super-resolution imaging and metabolic assays. (B) Representative confocal imaging and 3D renderings of 2D monolayers and 3D organoids from the PPN and PBM subtypes. Note differences in cytoplasmic surface area and organoid volume. (C) Heat map showing k-means clustering of proteins varying in PPN and PBM cells and/or in 2D and 3D conditions. Mean cluster patterns are shown across conditions as protein abundance z-scores. (D) Enrichment of Gene Ontology (GO) biological process terms for proteins within each k-means cluster in (C).

Prior to comparing the HGSC models in 2D and 3D, we assessed how they differed from parental cells. As expected, organoids of the PPN and PBM proteomes showed clear separation from parental *Trp53^-/-^* (P) by principal components analysis (Figure S1B, Supplementary Table 1). Genotype-dependent comparisons also revealed broad differences between PPN and P and PBM and P, respectively, including enrichment of programs related to mitotic progression, intracellular lipid transport, lipid catabolism, inflammatory/lipopolysaccharide response, and folate/amino-acid and cell-substrate regulatory pathway (Figure S1C, Supplementary Table 2).

Next, we compared the PBM and PPN proteomes when these lines were grown in 2D versus 3D conditions. We performed k-means clustering across PBM and PPN in monolayer and organoid conditions grown in their respective 2D and 3D media. We resolved six abundance-pattern clusters (Figure 1C, Supplementary Table 3). Cluster 1 was enriched in proteins elevated in PPN compared with PBM cells under both 2D and 3D conditions, with the 3D PPN condition further exhibiting higher abundances over the 2D condition. Gene Ontology (GO) analysis revealed enrichment for vesicle transport, lipid-related processes (metabolism, biosynthesis, localization), mitochondrial organization, Ca²⁺ homeostasis, membrane potential and fission, and microtubule polymerization and organization. This combination suggests that PPN cells may be predisposed toward reorganizing intracellular transport, membrane dynamics, and organelle communication. By contrast, Cluster 2 was enriched in PBM cells (over PPN) to a similar extent in both 2D and 3D (Figure 1D). Processes associated with this cluster included mitochondrial gene expression and translation, tRNA metabolism/modification, aminoacylation for translation, immune response, and DNA damage. In agreement, comparison of PPN and PBM organoids showed a broader set of underrepresented antiviral and innate immune-associated modules (Figure S1D, Supplementary Table 4), consistent with PPN having a lower basal abundance of proteins encoded by interferon-stimulated genes (Figure S1E). Cluster 3 captured an organoid-associated oxidative metabolic program (Figure 1D). This cluster was enriched in both organoid lines relative to monolayers, with the strongest increase in PBM, and was comparatively reduced in both monolayer conditions. It was characterized by oxidative phosphorylation, respiration, electron transport chain, ATP biosynthesis, and fatty acid metabolism/β-oxidation/lipid oxidation terms. Given this observation, it is possible that organoid growth favors or imposes mitochondrial oxidative metabolism and respiration relative to monolayer growth, irrespective of ovarian cancer genotype. Cluster 4 was most prominent under 2D growth conditions for both genotypes. Enriched GO terms for this cluster included ribonucleoprotein complex biogenesis, RNA processing and splicing, transcription, gene expression and translation, mitotic cell cycle, chromosome organization, and histone modification (Figure 1D). Cluster 5 showed elevated proteins in PPN monolayers relating to DNA metabolism and replication-associated programs, whereas cluster 6 highlighted elevated proteins in PBM monolayers involved in cell projection assembly, actin polymerization and intracellular transport, as well as RAS/GTPase-associated signaling.

Collectively, the clusters separate into programs that are PPN-, PBM-, organoid-, or monolayer-enriched, pointing to how genotype and growth format jointly partition the proteome. The reciprocal GO terms of Clusters 2 and 4 support the notion that 3D growth shifts cells away from proliferation-linked biosynthesis and toward oxidative and catabolic metabolism. The combination of Clusters 1 and 3 suggest that phenotypes in PPN might arise from the intersection of an organelle communication program and a shared oxidative organoid metabolic state.

### Media-controlled protein charting distinguishes shared and format-specific 3D states

To control for media-dependent proteomic features, we expanded the 2D monolayer versus 3D organoid analysis to include monolayers of each HGSC genotype cultured in both 2D and 3D media conditions (Figure 2A). Across PBM and PPN backgrounds, a principal components analysis assessed protein profiles upon culturing in 2D media (monolayers) or 3D media (monolayers and organoids). Organoids separated from monolayers, while the same growth format also shifted according to whether cells were grown in 2D or 3D media, indicating that the proteome is structured by both growth format and environmental media context. Upon growing 2D monolayers and organoids both in 3D media, differential enrichment mapping showed that proteins underrepresented in organoids were associated with cell cycle, mitotic cell cycle, ribosome biogenesis, rRNA metabolism, and microtubule/cytoskeleton organization (Figure 2B, Figure S2A, Supplementary Tables 5-9). By contrast, proteins overrepresented in organoids were enriched for lipid biosynthesis, fatty acid β-oxidation, ATP synthesis, oxidative phosphorylation, and extracellular matrix (Figure 2B, Figure S2A). When both monolayers and organoids are grown in 3D media to control for media-dependent changes, these results support that organoid growth is associated with a shift away from a proliferative phenotype with ribosomal biosynthesis, towards oxidative and lipid-linked metabolism.

**Figure 2.**
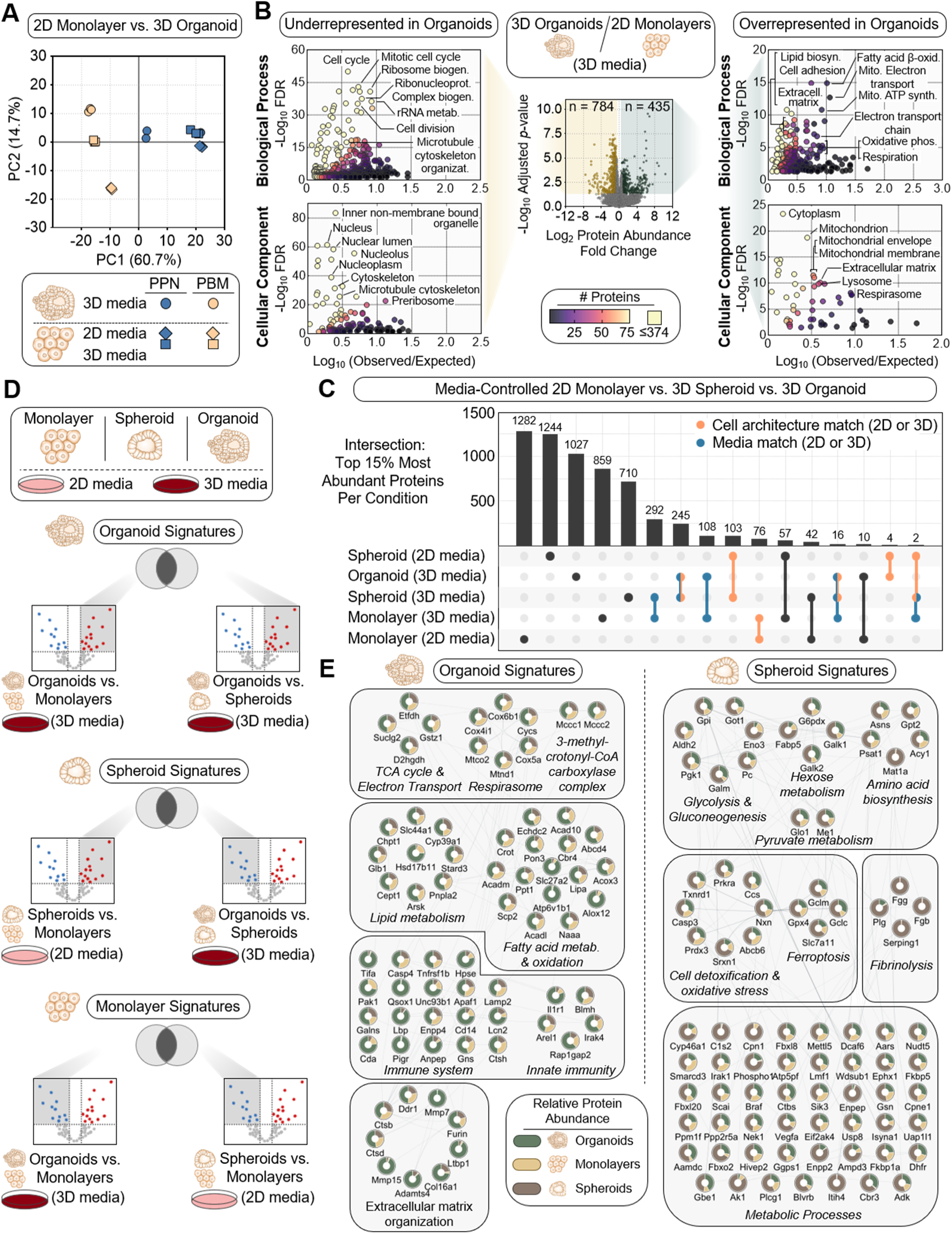
Media-controlled protein charting distinguishes shared and format-specific 3D states. (A) Principal Components Analysis of 2D monolayer and 3D organoid cultures in HGSC lines (*Trp53^-/-^;Pten^-/-^;Nf1^-/-^* (PPN) and *Trp53^-/-^;Brca1^+/-^;Myc^OE^* (PBM)), grown under 2D and 3D media conditions. (B) Differential enrichment analysis showing proteins under-(left) or over- (right) represented in 3D organoids relative to 2D monolayers grown in 3D media. Representative GO biological process and cellular component terms are shown. (C) UpSet plot showing intersection between the top 15% most abundant proteins per condition in 2D monolayers, 3D spheroids, and 3D organoids. Matches (shared proteins) driven by cell architecture (2D or 3D) and by media (2D or 3D) are highlighted. (D) Schematic showing conditions used to establish organoid and spheroid proteomic signatures. (E) Protein-protein interaction networks of organoid-enriched (left) and spheroid-enriched (right) protein signatures from (F), annotated by using the STRING database. Nodes display relative abundances in monolayers, spheroids, and organoids. Edges represent STRING functional protein interactions.

Next, we extended our analysis to include 3D spheroids grown in both 2D and 3D media. Spheroids are particularly useful for modeling a form of 3D growth because they establish cell-cell contacts, nutrient and oxygen gradients, and metabolic stress states that are not well captured in 2D monolayers^2, 8, 25^. Unlike organoids, they can also be generated in the absence of matrix embedment. Thus, including spheroids in the same matched framework allowed us to determine whether 3D-associated proteomic features are shared across tissue-like formats or are specific to ECM-embedded organoid growth. To compare major shared versus specific proteins in monolayers, spheroids, and organoids, respectively, we focused on the top 15% most abundant proteins per condition (Figure 2C). An UpSet plot revealed a large number of condition-specific proteins. However, there were also significant numbers of proteins whose expression was shared based on cell architecture (2D monolayers, or 3D organoids and spheroids) or when cultured in the same media (2D, or 3D media). There was also a shared fraction among 3D states (spheroid and organoid), consistent with a core 3D program. Pearson correlation analysis reinforced a within-condition clustering, together with separation between monolayer, spheroid, and organoid conditions (Figure S2B). Rank-ordered abundance plots further demonstrated a range of increased and decreased protein abundances across all intensity levels, reflecting broad and distributed state-level shifts (Figure S2C). These findings indicate that both architecture and media contribute to the proteomic state in a non-redundant manner. Some protein sets are shared across 3D conditions, whereas others are specific signatures to organoids, spheroids, or particular architecture–media combinations.

To determine these cell architecture signature profiles, we used a differential comparison scheme with multiple pairwise contrasts (Figure 2D, Figure S3). For example, organoid-enriched proteins were defined by comparing organoids versus monolayers and organoids versus spheroids (both grown in 3D media), whereas spheroid-enriched proteins were defined by overlap between spheroids versus monolayers (2D media) and organoids versus spheroids (3D media). GO analysis showed that the organoid-enriched proteins resolve into modules that couple mitochondrial respiration, lipid utilization, lysosomal remodeling, and extracellular matrix organization, consistent with a tissue-like state sustained by oxidative and matrix-interactive metabolism (Figure 2E, left, Supplementary Table 5). By contrast, spheroid-enriched proteins include glycolysis, pyruvate and hexose metabolism, ferroptosis-related factors, and oxidative detoxification, consistent with cells adapting to redox challenge (Figure 2E, right, Supplementary Table 5).

Taken together, 3D culture formats derived from the same cells can yield systematically different biological readouts. Organoids and spheroids share a core 3D program, but diverge into metabolically distinct states. The observations indicate that organoids adopt a matrix-supported oxidative and remodeling state, whereas spheroids preferentially engage a stress-buffering, glycolytic survival state. Organoid effects may reflect a combination of architecture and media context, rather than architecture alone. This interpretation is supported by our monolayer-enriched protein analysis, in which signature proteins were organized into DNA replication, checkpoint, mitosis, and ribonucleoprotein biogenesis modules (Figure S3A). Proteins enriched in spheroids over monolayers depended specifically on 3D media highlighted consistent architecture-by-media biology (Figure S3B).

### A microtubule-enriched Trp53^-/-^;Pten^−/−^;Nf1^-/-^(PPN) state coincides with paclitaxel hypersensitivity

Having established cell architecture-specific states, we next asked whether ovarian cancer genotype-specific biases within these states could reveal mechanistically informative or actionable biology. Microscopy showed differences in cytoplasmic surface area (Figure 1B) and Cluster 1 analysis showed that PPN, versus PBM, are enriched in proteins with roles in microtubule polymerization and organization (Figure 1C). These observations suggested that PPN might have a different cytoskeletal organization than PBM. As we reported earlier^15^, PPN monolayer cells showed greater sensitivity to paclitaxel than PBM in viability assays (Figure 3A). DAPI imaging revealed correspondingly stronger paclitaxel-associated micronucleation in PPN relative to PBM cells (Figure 3B).

**Figure 3.**
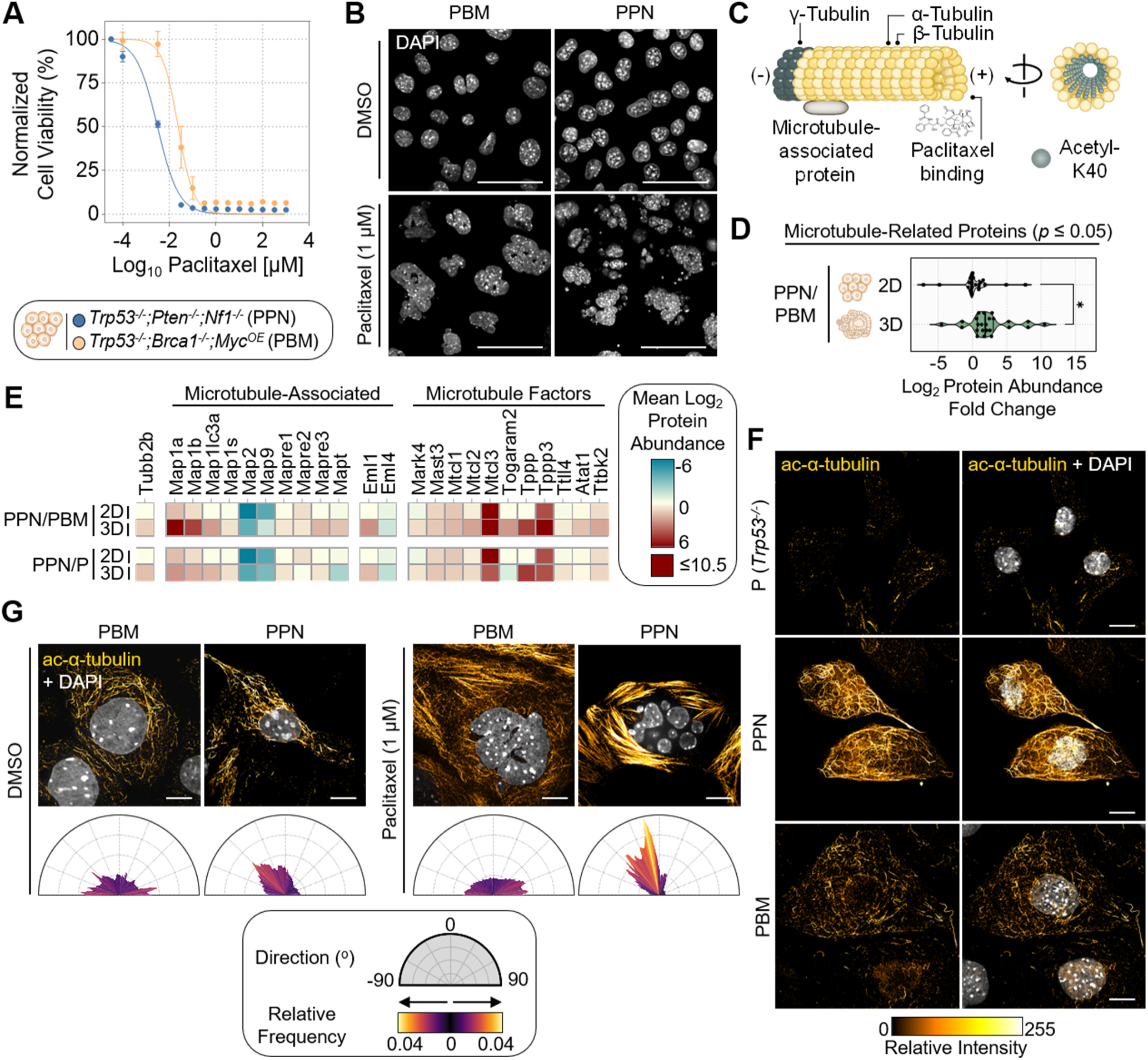
A microtubule-enriched PPN cell state coincides with paclitaxel hypersensitivity. (A) Paclitaxel dose-response curves in HGSC monolayer cells in 2D media (*Trp53^-/-^;Pten^-/-^;Nf1^-/-^*(PPN) and *Trp53^-/-^;Brca1^+/-^;Myc^OE^* (PBM)). Normalized cell viability was calculated relative to control-treated cells (0.01% DMSO) and measured after treatment for 5 days. (B) Representative DAPI images of HGSC PBM and PPN cells following DMSO vehicle or paclitaxel treatment for 20 h. (C) Schematic of microtubule and paclitaxel binding. Acetylated α-tubulin at K40 marks stable microtubules. (D) Comparison of microtubule-related protein abundances exhibiting *p* ≤ 0.05 between PPN and PBM across 2D monolayer and 3D organoid conditions. Median (thick black line) and quartiles (thin black line) are shown. Statistical significance was determined using Welch’s t-test (\**P* < 0.05). (E) Heat maps of microtubule-associated and microtubule regulatory protein abundances from (D) across PBM, PPN, and parental (*Trp53^-/-^* (P)) states. (F) Representative confocal imaging of acetylated α-tubulin (K40) across parental P, PBM, and PPN cells, showing differences in the abundance and organization of stable microtubules. (G) Representative confocal imaging of acetylated α-tubulin (K40) of PBM and PPN cells upon DMSO vehicle or paclitaxel treatment for 20 h. Directionality plots quantify the relative orientation and frequency of microtubule organization in the representative cells.

We examined whether the paclitaxel-induced differences between PPN and PBM cells might be explained by differences in microtubule biology, or relative sensitivity to cell death in response to microtubule disruption. Paclitaxel binds β-tubulin within polymerized microtubules (Figure 3C), suppressing dynamic instability, stabilizing bundled microtubule structures, and ultimately perturbing spindle function and chromosome segregation^26, 27^. Microtubule-related proteins were elevated in PPN relative to PBM or P, particularly in 3D conditions (Figure 3D), including structural tubulins, microtubule-associated proteins (MAPs), and microtubule-linked factors (Figure 3E). These observations are consistent with prior reports that NF1 associates with microtubules^28, 29^ and prior observation that NF1 is responsible for the paclitaxel hypersnsitivity^15^. Using microscopy, we observed that PPN cells display a more prominent acetylated α-tubulin network, indicative of stable microtubules (Figure 3F). Under vehicle and paclitaxel treatment, PPN exhibited pronounced bundled and directionally aligned acetylated microtubule structures (Figure 3G). Directionality analysis showed a more organized, anisotropic network upon treatment. Taken together, these results support a model in which a PPN-enriched microtubule state is associated with increased susceptibility to paclitaxel-induced cytoskeletal disruption.

### Mitochondria-ER mediated lipid droplet clustering is associated with the PPN state

Cluster 1 suggested that PPN cells have distinct lipid metabolism, membrane dynamics, and organelle communication (Figure 1C-D). Cellular component analysis of the PPN-enriched lipid proteins showed that they were strongly biased toward ER membrane, mitochondria-associated ER membrane, mitochondrial membrane compartments, and lipid droplets (Figure 4A, Supplementary Table 10). These observations suggested that the PPN state has an altered lipid program with enhanced mitochondria-ER contacts. Indeed, BODIPY staining of live cells showed that, relative to parental and PBM, PPN cells exhibited both an increase in lipid signal and pronounced lipid droplet clustering (Figure 4B, S4A-B).

**Figure 4.**
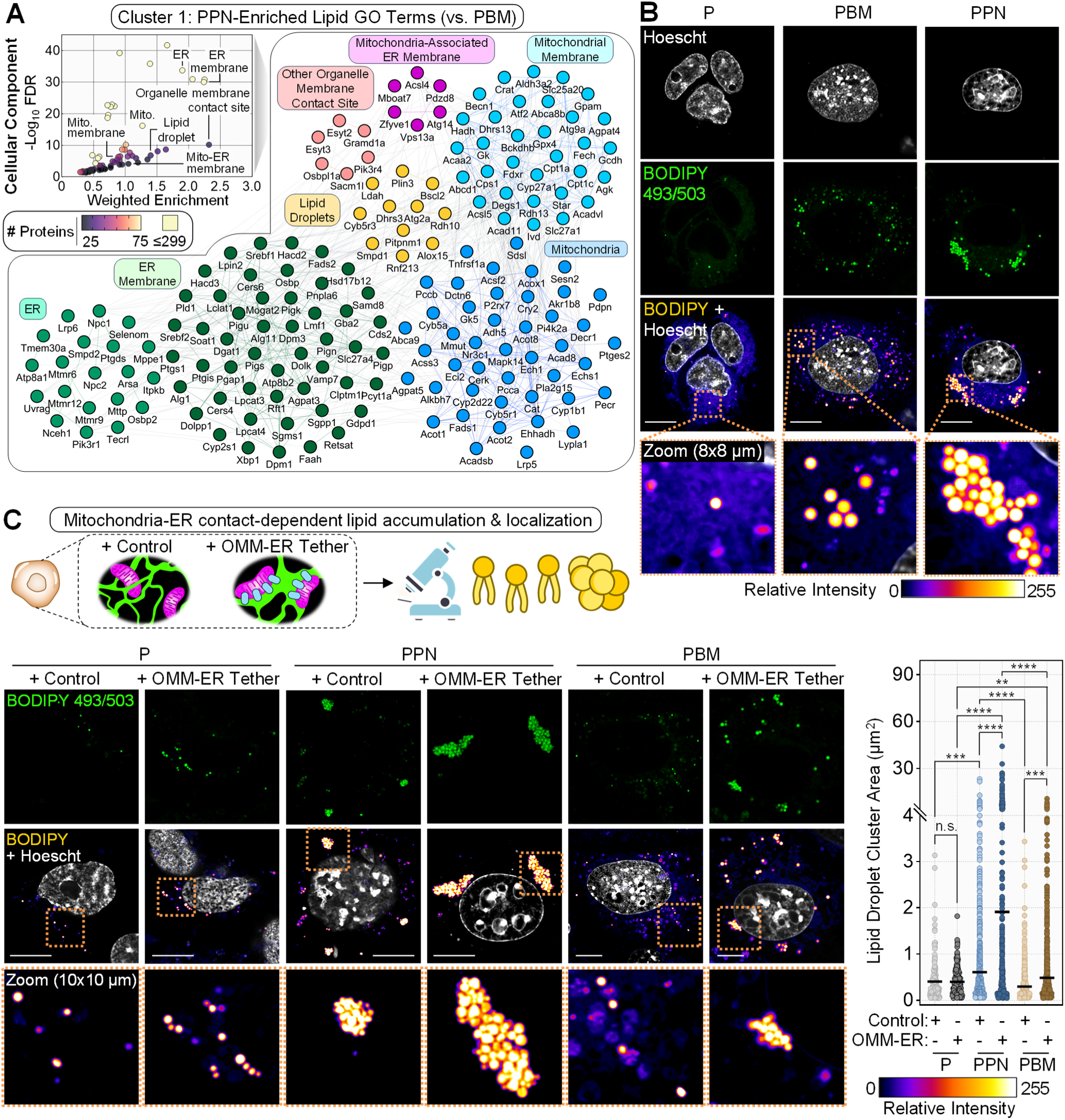
Mitochondria-ER mediated lipid droplet clustering is associated with the PPN state. (A) Cellular component enrichment and protein-protein interactions of lipid-associated proteins elevated in *Trp53^-/-^;Pten^-/-^;Nf1^-/-^* (PPN) relative to *Trp53^-/-^;Brca1^+/-^;Myc^OE^* (PBM) cells (Cluster 1 in Figure 1C-D). (B) Representative confocal imaging upon BODIPY 493/503 staining across parental (*Trp53^-/-^*(P)), PBM, and PPN cells showing lipid droplet organization. (C) Representative confocal images of BODIPY 493/503 staining in P, PPN, and PBM cells upon expressing a control or outer mitochondrial membrane-ER tether construct. Quantification of lipid droplet cluster area by two-way ANOVA and Tukey post hoc test (n.s., not significant; \*\**P* < 0.01; \*\*\**P* < 0.001; \*\*\*\**P* < 0.0001). Solid black line indicates the mean.

To test whether enhanced mitochondria-ER tethering drives clustered lipid droplets, we expressed an outer mitochondrial membrane-ER tether (Figure 4C). The tether had little effect in the parental background, but increased lipid droplet clustering in both transformed genotypes, with the strongest effect in PPN. The context dependence of this effect indicates that enhanced tethering can reorganize lipids in transformed states that are already primed for mitochondria-ER output, as opposed to being universally sufficient in every intracellular background.

### PPN cells are also enriched for mitochondrial-ER remodeling

PPN-enriched proteins in cluster 1 converged on processes that rely on spatial coordination between organelles, e.g., microtubule organization with lipid metabolism, mitochondrial organization, Ca²⁺ homeostasis, and membrane dynamics (Figure 1C-D). We therefore asked whether PPN cells show differences in inter-organelle communication, including between the mitochondria and ER. We sought to quantify membrane contact site proteins (MCSs), which physically and functionally span inter-organelle contact interfaces. The detection of MCSs can be difficult due to their frequent low endogenous levels and membrane-bound characteristics, impeding their quantification in conventional non-targeted mass spectrometry (MS) approaches. We adapted our reported targeted MS approach based on parallel reaction monitoring^30^ to simultaneously quantify MCSs spanning major inter-organelle interfaces (Figure 5A, Supplementary Table 11). MCS levels were elevated in PPN organoids relative to PBM organoids, with the strongest increase in mitochondria-ER contacts (Figure 5A-B). Relative to monolayers, organoid growth further increased mitochondria-ER MCSs in PPN cells and also revealed increases in ER-endolysosome and outer mitochondrial membrane-inner mitochondrial membrane (OMM-IMM) contact-associated proteins.

**Figure 5.**
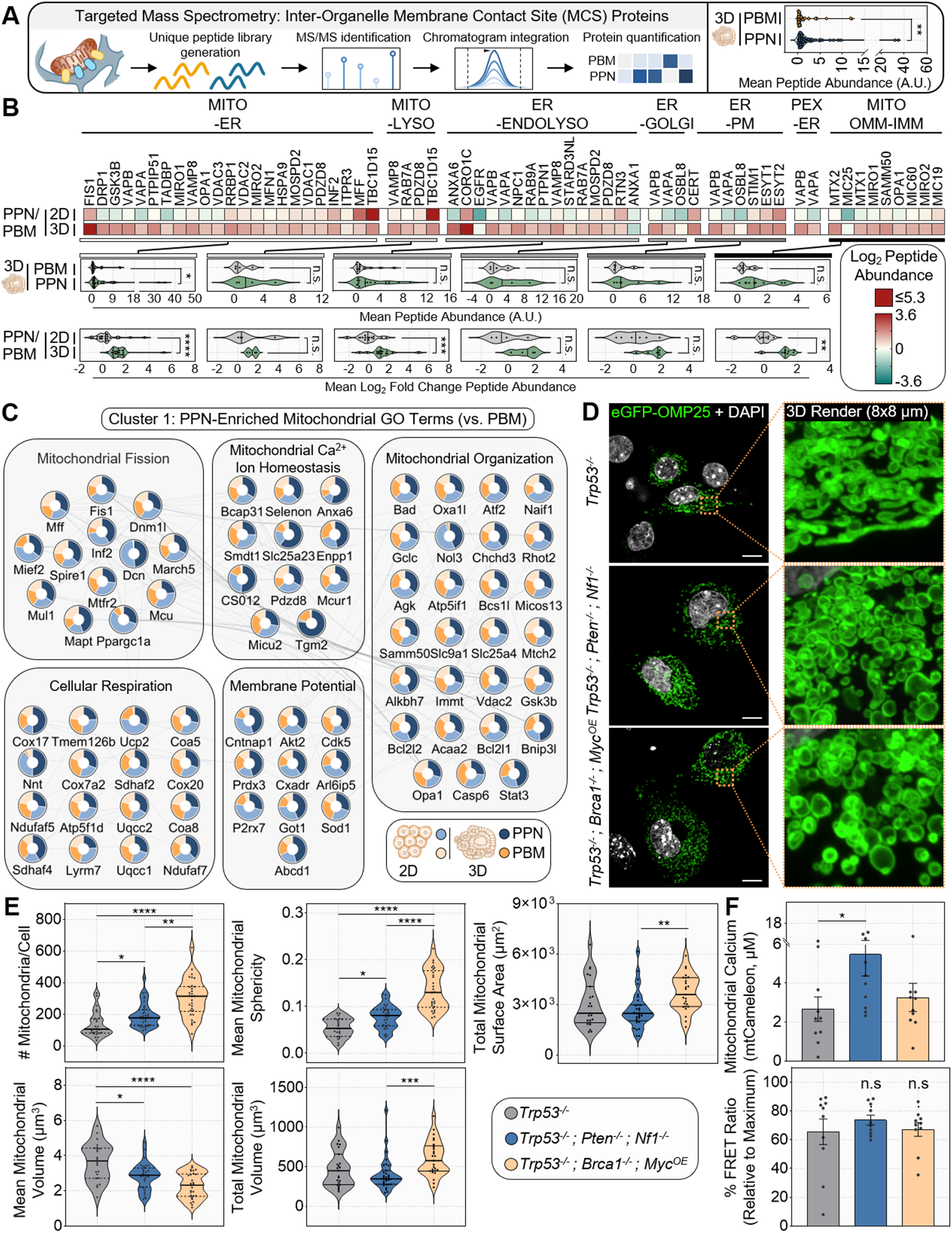
PPN cells show evidence of mitochondrial-ER remodeling. (A) Workflow for targeted mass spectrometry using parallel reaction monitoring to quantify functional inter-organelle membrane contact site (MCS) protein abundances in the HGSC cell models. Proteotypic peptides were monitored and a summary violin plot of total MCS peptide abundances between *Trp53^-/-^;Pten^-/-^;Nf1^-/-^*(PPN) relative to *Trp53^-/-^;Brca1^+/-^;Myc^OE^* (PBM) organoids is shown. Statistical significance was determined with a Mann-Whitney test (\*\**P* < 0.01). (B) Heat maps and violin plots of quantified MCS protein abundances in HGSC cells grown as 2D monolayers and 3D organoids. Mean peptide abundances are depicted. MCS proteins were categorized as mediating functional contacts between mitochondria-endoplasmic reticulum (ER), mitochondria-lysosome (lyso), ER-endolysosome (endolyso), ER-Golgi, ER-plasma membrane (PM), peroxisome (PEX)-ER, and outer mitochondrial membrane - inner mitochondrial membrane (mito OMM-IMM). Median (black line) is shown. Statistical significance was determined with a Mann-Whitney test (n.s., not significant; \**P* < 0.05; \*\**P* < 0.01; \*\*\**P* <0.001; \*\*\*\**P* < 0.0001). (C) Protein-protein interaction network of mitochondrial GO terms enriched in PPN cells (Cluster 1 in Figure 1C-D). Proteins were annotated using the STRING database. Nodes display relative abundances on PPN and PBM samples (2D monolayers and 3D organoids). Edges represent STRING functional protein interactions. (D) Representative confocal imaging and 3D rendering of mitochondrial architecture upon expression of eGFP-OMP25 in parental (*Trp53^-/-^* (P)), PBM, and PPN cells. (E) Quantification of mitochondrial number per cell, sphericity, surface area, mean volume, and total mitochondrial volume in parental P, PBM, and PPN cells. Median (solid black line) and quartiles (dotted black line) are shown. Statistical significance was determined using the Kruskal-Wallis test (\**P* < 0.05; \*\**P* < 0.01; \*\*\**P* <0.001; \*\*\*\**P* < 0.0001). (F) Quantification of mitochondrial calcium levels and % FRET ratio using expression of mtCameleon in cells. Mean with SEM error bars are shown. Statistical significance was determined using the Kruskal-Wallis test (n.s., not significant; \**P* < 0.05).

Because mitochondria-ER and OMM-IMM interfaces are major determinants of mitochondrial morphology, calcium exchange, and respiratory organization^31–33^, we next asked whether the MCS-rich PPN state coincides with mitochondrial remodeling. Consistent with this, mitochondrial GO terms elevated in PPN relative to PBM cells included mitochondrial fission, mitochondrial calcium ion homeostasis, mitochondrial organization, cellular respiration, and membrane potential (Figure 5C). The mitochondrial fission module included FIS1, DNM1L, MIEF2, INF2, MUL1, and MARCHF5, a combination suggestive of enhanced mitochondrial division and remodeling. This observation was supported by specific proteins in the targeted MS dataset, including fission-and remodeling-associated factors (e.g., FIS1, DRP1, MFF) and related mitochondrial membrane organizers. Overall, these observations suggest that the MCS-enriched PPN state is accompanied by a shift in mitochondrial architecture. The mitochondrial calcium ion homeostasis module contained MCU, MICU2, SMDT1, SLC25A23, CISD2, and BCAP31/SELENON-associated regulators, all of which raise the possibility that calcium exchange between mitochondria and neighboring membranes is altered in PPN cells. Finally, the mitochondrial organization and OMM–IMM contact modules included proteins (e.g., CHCHD3, MICOS13, SAMM50, MTCH2, OXA1L) consistent with remodeling of inner membrane architecture, cristae organization, and communication between intramitochondrial compartments. These features, together with respiratory assembly factors (e.g., COX17, TMEM126B, COX20, UQCC1/2, ATP5F1D, SDHAF2/4, NDUFAF7), further suggested that the mitochondria in PPN cells could be organized into a metabolically distinct state.

Given the proteomic-based prediction of heightened mitochondrial fragmentation or fission in PPN cells, we used confocal imaging to visualize mitochondria in parental P, PBM, and PPN cells. PPN cells displayed increased numbers of mitochondria, with reduced mean mitochondrial volume and increased sphericity (Figure 5D and Figure S5). We also assessed mitochondrial Ca^2+^ levels by expressing the Ca^2+^ sensor, mtCameleon^34, 35^. mtCameleon localizes to mitochondria and exhibits elevated fluorescence upon binding to Ca^2+^. mtCameleon measurements showed that PPN cells exhibited increased mitochondrial calcium relative to PBM and the parental cells, whereas the maximal normalized FRET response was altered less (Figure 5F). This selective increase in mitochondrial calcium load is consistent with the elevation of calcium-handling machinery and with the broader enrichment of mitochondria-ER MCS proteins in PPN. Together, these results support a model in which PPN cells are characterized by increased mitochondria-ER contacts, mitochondrial fission, and mitochondrial calcium. These characteristics could alter the architecture and stability of mitochondria-ER interfaces in cells and thereby influence respiratory behavior.

### PPN exhibits mitochondria-ER encapsulations associated with increased cellular respiration and cytomegalovirus permissiveness

To determine whether the mitochondria-ER MCS profiles in PPN and PBM correspond to altered organelle behavior in living cells, we examined mitochondria-ER contact dynamics by live-cell confocal imaging (Figure 6A, Figure 6B). We monitored the extent to which mitochondria and ER can undergo dynamic tubule crossing, or exhibit more stable mitochondria-ER interactions. Representative images from PBM and PPN cells showed that the latter more frequently exhibited mitochondria-ER contacts (Figure 6A). Time-lapse imaging, revealed that in PBM cells mitochondria–ER contacts often appeared as transient crossings or partial alignments (Figure 6B). By contrast, PPN cells exhibited stable, encapsulation-like arrangements that persisted over time. Quantification confirmed a shift away from dynamic and no-contact phenotypes toward stable mitochondria-ER contacts in PPN relative to PBM and parental cells (Figure 6C). The stability of the contacts was reminiscent of mitochondria-ER encapsulations (MENCs), which we previously described to facilitate protection of fragmented mitochondria from mitophagy and increased oxidative phosphorylation in the context of infection with human cytomegalovirus (HCMV), which is associated with oncomodulation^30, 36^.

**Figure 6.**
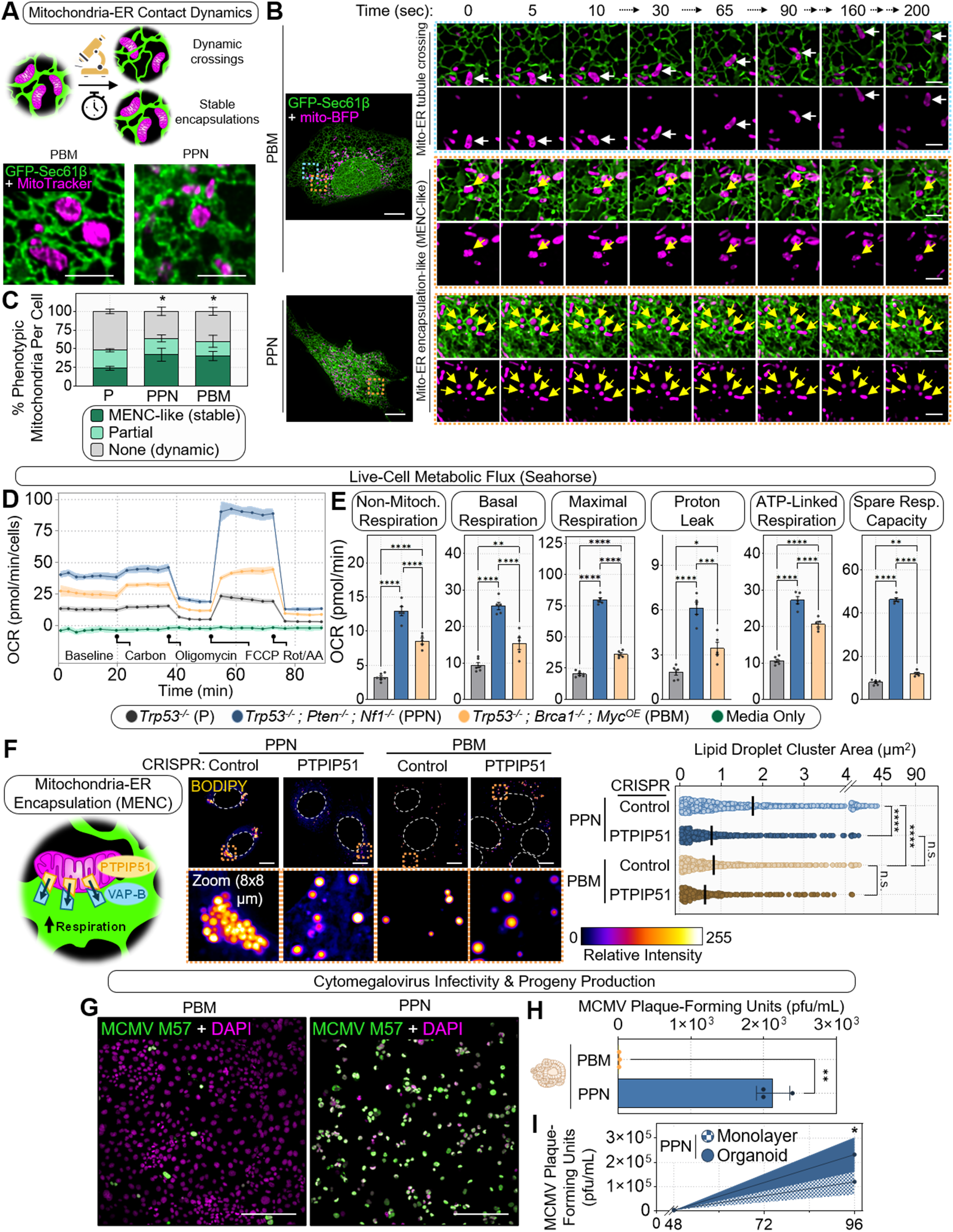
PPN cells show altered mitochondrial dynamics associated with increased cellular respiration and cytomegalovirus permissiveness. (A) Schematic showing overview of mitochondria-ER contact dynamics, including transient crossings and stable, asymmetric mitochondria-ER encapsulations (maintenance of mitochondria-ER contact ≥120 s). Representative confocal image of mitochondria (MitoTracker) and ER (GFP-Sec61β) in HGSC cells (*Trp53^-/-^;Pten^-/-^;Nf1^-/-^*(PPN) and *Trp53^-/-^;Brca1^+/-^;Myc^OE^* (PBM)). (B) Representative live-cell confocal imaging illustrating mitochondria-ER contact behavior: dynamic mitochondria-ER crossing in PBM cells and stable, asymmetric mitochondria-ER encapsulations in PBM and PPN cells are shown. Scale bar (whole cell): 10 µm; scale bar (zoom inset): 2 µm. (C) Quantification of mitochondria-ER contact state frequencies in parental P, PBM, and PPN cells (stable, asymmetric mitochondria-ER encapsulations: maintenance of contact ≥120 s; partial: maintenance of contact ≥45 s, < 120 s). Mean with SD error bars are shown. Statistical significance was determined using a Welch’s t-test (\**P* < 0.05). (D-E) Oxygen consumption rate (OCR) and associated respiratory parameters determined by using a live-cell Seahorse assay. Summary respiratory parameters include non-mitochondrial respiration, basal respiration, maximal respiration, proton leak, ATP-linked respiration, and spare respiratory capacity. Mean with SEM error bars are shown. Statistical significance was determined using one-way ANOVA and Tukey post hoc test (\**P* < 0.05; \*\**P* < 0.01; \*\*\**P* <0.001; \*\*\*\**P* < 0.0001). (F) Schematic showing mitochondria-ER encapsulation and known MCS proteins PTPIP51 and VAP-B, which relocalize to MENCs. Representative confocal imaging of BODIPY 493/503 staining in PPN and PBM cells deleted for *Ptpip51* (Control and PTPIP51). Quantification of lipid droplet cluster area is shown at right. Scale bar: 10 µm. Statistical significance was determined using two-way ANOVA and Tukey post hoc test (n.s., not significant; \*\*\*\**P* < 0.0001). Solid black line indicates the mean. (G) Representative confocal imaging of PBM and PPN cells infected with murine cytomegalovirus (MCMV, 48 h post-infection, multiplicity of infection of 1). MCMV M57 is the viral major single-stranded DNA binding protein. (H) Quantification of MCMV infectious progeny production as plaque-forming units in PBM and PPN organoids (72 h post-infection, multiplicity of infection of 1). Mean and SD error are shown. Statistical significance was determined using Welch’s t-test (\*\**P* < 0.01). (I) Quantification of MCMV infection progeny production as plaque-forming units in PPN monolayers and organoids (multiplicity of infection of 0.05). Mean and SD error are shown. Statistical significance was determined using Welch’s t-test (\**P* < 0.05).

Because we and others have (1) demonstrated that mitochondria-ER contacts, and MENCs in particular, regulate metabolism, substrate flux, and calcium transfer, and (2) PPN cells have altered mitochondria architecture (Figure 5), we next measured bioenergetic behavior. Live-cell metabolic flux analysis using a Seahorse instrument showed that PPN cells displayed the highest oxygen consumption and extracellular acidification rates (Figure 6C-D, Figure S6A-B). Respiratory parameters confirmed that PPN had elevated non-mitochondrial respiration, basal respiration, maximal respiration, proton leak, ATP-linked respiration, and spare respiratory capacity relative to PBM and the parental cells (Figure 6C-D, Figure S6B). These data support the notion that stable mitochondria-ER contact organization is associated with a high-flux metabolic state, characterized by both elevated mitochondrial respiration and increased glycolytic acidification. The greater spare respiratory capacity is consistent with the increased mitochondrial number and remodeled mitochondrial architecture observed in PPN (Figure 5), suggesting that the fission-biased mitochondrial state may expand respiratory reserve.

To test whether endogenous MENC-associated tethering machinery contributes to downstream spatial lipid remodeling (e.g. lipid droplet clustering), we disrupted the gene encoding the mitochondria-ER contact protein PTPIP51 by CRISPR/Cas9 gene editing (Figure 6E; Figure S6C). Previous work demonstrated that PTPIP51, along with VAP-B, relocalizes to MENCs during HCMV infection, becomes increasingly enriched at MENCs as infection progresses, and is required for the infection-associated increase in mitochondrial respiration^30, 36^. *PTPIP51* deletion reduced the size of the clustered lipid droplets most strongly in PPN cells. Thus, both mitochondria-ER tethering and disruption support a connection between mitochondria-ER communication and lipid droplet organization in the transformed states.

CMV infection is known to exploit mitochondria-ER contact dynamics and form MENCs, promoting lipid metabolism and mitochondrial bioenergetics. The features of PPN cells thus far suggested that they might show increased CMV permissiveness. PPN cells also expressed lower levels of proteins encoded by antiviral interferon-stimulated genes PPN also expressed lower levels of antiviral interferon-stimulated genes (Figure S1C-D). Consistent with our predictions, imaging of murine CMV (MCMV) infection imaging (Figure 6F), and plaque assays confirmed increased susceptibility of PPN compared with PBM cells (Figure 6G). The output of infectious virus was also higher in PPN than in PBM organoids, PPN organoid cultures also yielded greater infectious output than monolayers (Figure 6H), linking the organoid state itself to enhanced permissiveness. Together, these data support a model in which stable mitochondria-ER contacts, increased respiratory reserve, elevated glycolytic acidification, lipid droplet clustering, and a diminished basal antiviral profile facilitates a cellular environment unusually permissive for cytomegalovirus infection.

## Discussion

Three-dimensional culture systems are increasingly used across cell biology, cancer, development, and host-pathogen biology. Yet, the molecular differences amongst monolayers, spheroids, and organoids remain insufficiently resolved. Here, we show that monolayer, spheroid, and organoid growth conditions evoke distinct proteomes and that these states can have functional consequences. By separating the architecture from media context, we provide a unique, reusable dataset that distinguishes shared 3D programs from format-specific states. We demonstrate the utility of predictions generated by this resource by showing how HGSC cells of defined genotype, particularly when cultured in 3D, link ER–mitochondria contacts, mitochondrial respiration, lipid droplet clustering, and cytomegalovirus permissiveness.

Our results show that organoids and spheroids are not interchangeable 3D systems. We find that organoids exhibit oxidative, lipid-linked, lysosomal, and ECM-remodeling programs, whereas the same cells cultured as spheroids are enriched for glycolysis, redox buffering, and ferroptosis-protective pathways. These differences likely reflect the distinct environment and constraints imposed by each format (i.e., matrix-supported, tissue-like organization in organoids versus diffusion-limited, anoikis-inducing stress adaptation in spheroids). More generally, these data emphasize that culture format is a key determinant of cellular state and provide an explanation for why different 3D systems can yield different conclusions when used to model the same tissue. Prior work comparing 2D and 3D cultures has emphasized differences in cell-cell and cell-matrix interactions, nutrient gradients, and metabolic behavior^1, 2, 4, 7, 37–39^. Our data extend complement and extend these studies by providing a proteome-wide resource that distinguishes shared 3D biology from format-specific programs. A second principle emerging from our study is that architecture and media jointly shape the proteome. We observed not only strong main effects of growth format, but also proteins and pathways whose abundance patterns depended on the combination of architecture and media. Consequently, previously observed differences in cell behavior in 2D vs 3D likely reflect a combination of geometry, matrix, and median conditions, rather than dimensionality alone. Our data therefore provide a practical framework for benchmarking 2D and 3D models and for interpreting discrepant results across experimental systems. Our data suggest that monolayers, spheroids, and organoids model different tumor-relevant states. For example, HGSC often disseminates through the peritoneal cavity as multicellular spheroids in ascites, and these spheroids can attach to and clear mesothelial surfaces during implantation^40, 41^. Thus, spheroids may be particularly useful for modeling anchorage-independent aggregate survival and early peritoneal dissemination, whereas ECM-embedded organoids may better capture matrix-engaged tumor deposits and tissue-like metabolic states. Monolayers are useful for scalable perturbation and substrate-attached proliferative states, yet do not necessarily reproduce the architecture or diffusion constraints of 3D growth. These differences may be impact drug responses and it might be necessary to test all source-matched models.

The mechanistic findings of the study emerged from an unexpected convergence. Proteomic clustering first highlighted a microtubule-enriched PPN cells, which correlated with paclitaxel hypersensitivity. We found that PPN cells were also enriched for MCS components, mitochondrial remodeling, elevated respiration, and lipid droplet clustering. Together, targeted MS, live-cell imaging, calcium measurements, and perturbation experiments support a model in which contact-rich organelle organization stabilizes a respiratory cell state. Conceivably, one consequence of this state may be increased dependence on microtubule organization for maintaining organelle positioning and metabolic coordination, thereby enhancing vulnerability to paclitaxel despite the aggressive phenotype. Whether these are separate or related observations awaits future studies.

The differences in MCSs in PPN and PPM cells also provides an explanation for the lipid droplet clustering and viral permissiveness. In PPN cells, stable ER–mitochondria contacts correlate with elevated respiration, and clustered lipid droplets. Enhancement of mitochondria-ER tethering is sufficient to drive lipid droplet clustering in transformed backgrounds, whereas PTPIP51 loss suppresses clustering selectively in the most mitochondria-ER contact-enriched state. Therefore, we propose that one consequence of stable mitochondria-ER organization is the formation of spatially organized lipid domains that support oxidative metabolism. A similar observation has been made in lung adenocarcinomas exhibiting high oxidative phosphorylation^42^. Because cytomegalovirus exploits the host metabolic and secretory machinery and has been shown to remodel membrane contacts and organelles during infection, a mitochondria-ER rich, oxidative, lipid-organizing state with diminished basal antiviral tone would be predicted to favor infection^30, 36, 43–48^. Our infection data are consistent with this model.

Our study also has important limitations, particularly with respect to human pathobiology. We used genetically defined mouse cell systems, rather than patient derived cells, and the infection experiments tested MCMV rather than HCMV. In addition, we focused on two HGSC genotypes, which does not come close to representing the marked genetic heterogeneity in this disease. Nevertheless, we suspect that these are limitations of scope, not of principle. We established a proteomic framework for comparing 2D and 3D states, and further show that organoids can be connected coherently to organelle contacts, metabolism, therapy response, and viral permissiveness.

In summary, this study provides both a searchable proteomic reference and a mechanistic framework for many future studies. We find that architecture and media jointly structure the proteome; 3D states are metabolically and functionally distinct; and at least one HGSC genotype, when cultured in 3D, exhibits an mitochondria-ER contact dependent pathway linking lipids, and viral permissiveness. We anticipate that this framework will be useful in contexts where monolayers, spheroids, and organoids are used to model tissue-like biology.

## Experimental Procedures

### Cell and Organoid Culture

Murine P, PPN, and PBM monolayer cells and organoids were grown as previously described^15^. Organoids were thawed, seeded in Matrigel®, and cultured in 3D organoid growth medium, composed of: Ad+++ AdDMEM/F12 (Invitrogen); HEPES (Thermo Fisher, 100× diluted); and penicillin/streptomycin and GlutaMAX, each 100× diluted (Life Technologies), supplemented with B27 (Invitrogen, 50× diluted), N2 supplement (Thermo Fisher, 100× diluted), 1.25 mmol/L N-acetylcysteine (Sigma), 50 ng/mL EGF (Thermo Fisher), 500 ng/mL RSPO1 (PeproTech) or R-spondin-1 conditioned medium (25%–50%, v/v), WNT3a conditioned medium (0%–25%, v/v), 100 ng/mL Noggin (PeproTech), 50 ng/mL EGF, 10 ng/mL FGF10, 0.5 μmol/L A83-01 (Thermo Fisher), and 10 μmol/L Y-27632 (Sigma-Aldrich). Monolayers were derived from the organoids by dissociating organoids into single cells and plating on tissue culture-treated dishes. Spheroids were generated by plating monolayer cells at 2e6 cells/mL in ultra-low attachment microplates (Corning) for 24 h. Two-dimensional media contained DMEM with L-glutamine, 4.5g/L glucose and sodium pyruvate (Corning) supplemented with 3% fetal bovine serum albumin, 1x insulin-transferrin-selenium (Gibco), 50 ng/mL EGF, and penicillin/streptomycin.

### Transfection and CRISPR Knockout

To transfect cells, cells were grown to 70% confluency and media was refreshed to OptiMEM (Life Technologies). X-tremeGENE HP (Sigma Aldrich) transfection reagent was used in combination with the indicated DNA plasmids in a 1:3 ratio in OptiMEM and added dropwise to the plate. After 6 h incubation, cells were split into new dishes. PTPIP51 knock out cells were generated using two sequential rounds of transfection with TrueCut Cas9 protein V2 and CRISPRMAX Cas9 transfection reagent (Thermo Fisher) following the manufacturer’s protocols. TrueGuide Synthetic sgRNA (Control: Negative Control, non-targeting 1, sgRNA #1: CRISPR118552_SGM, sgRNA #2: CRISPR1038895_SGM) were used.

### Paclitaxel Sensitivity Assay

Paclitaxel sensitivity was determined in cells as previously described^15^. Monolayer cells derived from organoids were seeded into 96-well plates at 1,000 cells/well (day 0). Paclitaxel (Selleckchem, S1150) or 0.01% DMSO was added on the day following seeding (day 1). Media were refreshed and fresh drug was added on day 3. Cell viability was assessed on day 5 upon the addition of PrestoBlue HS and incubating for 30 minutes at 37°C. Fluorescence was measured using a Spectramax iD5 microplate reader (Molecular Devices). Results were normalized to the DMSO control condition and IC_50_ values were determined using GraphPad Prism.

### MCMV Progeny Production Assay

MCMV titers were determined by the TCID_50_ method. After the indicated cell pre-treatments, infections were performed at a multiplicity of infection of 1 (organoids) or 0.05 (monolayers). Viral inoculum was prepared by diluting the virus stock in media. At the indicated time post-infection, monolayer cells or organoids were scraped into the media and samples were freeze-thawed for three successive cycles using liquid nitrogen and a 37°C water bath. To remove cell debris, lysates were centrifuged at 20,000 ×g for 5 min at 4°C. Serial dilution of lysates in DMEM containing 2% v/v FBS and 1% v/v Pen/Strep was prepared to infect confluent monolayers of Vero cells. After 1 h viral adsorption at 37°C, viral inoculum was aspirated and replaced with 2% v/v FBS, 1% v/v Pen/Strep, and 1% methocel. Cell monolayers were incubated at 37°C until plaques were visible (∼3 days), washed in PBS, and fixed and stained with 50% v/v methanol, 1% w/v crystal violet for 15 min. Virus titers were calculated as plaque-forming units (PFU/mL).

### Immunoblot Analysis

Cells were lysed in 1x Laemmli buffer (62.5 mM Tris-HCl, pH 6.8, 2% w/v SDS, 10% v/v glycerol, 0.02% w/v bromophenol blue, 100 mM DTT), boiled at 95°C for 10 min, and run using SDS-PAGE. PVDF membrane was used to transfer proteins by the wet transfer method. 5% w/v milk in PBS was used to block membranes in 1x tris-buffered saline, and blots were incubated with primary and secondary antibodies. Immunofluorescence detection was performed to visualize protein bands using the LI-COR Odyssey CLx imaging system. The following antibodies were used: α-PTPIP51 (Santa Cruz Biotechnology; sc-166412) and α-tubulin (Sigma-Aldrich; T6199). Secondary antibodies were conjugated with Alexa Fluor fluorophore (Life Technologies).

### Immunofluorescence Imaging

All cells were seeded on 35 mm glass bottom microscope dishes (MatTek) pre-treated with fibronectin. A Nikon CSU W1 with Super Resolution by optical reassignment (SoRa) with a 60× Plan-Apo oil-immersion objective and a SoRa magnifier was used. Fixed immunofluorescence samples were fixed with 4% paraformaldehyde for 15 min at room temperature and permeabilized with 0.5% TritonX-100 for 10 min at room temperature. Cells were blocked in PBS containing 0.2% Tween-20 (PBS-T) with 2% goat serum and 2% bovine serum albumin. Samples were incubated in primary antibody for 1 h at room temperature, washed three times with PBS-T, and incubated in AlexaFluor secondary antibody at 1:2,000 and DAPI for 1 h at room temperature. The following antibodies were used: α-alpha tubulin (acetyl K40) (Abcam; ab179484) and α-MCMV M57 (Biorbyt; orb1101168). In live cell experiments, CellBrite® Steady 650 (Biotium; 30108) was used to stain the cell surface of monolayers and organoids and Hoescht (1 µg/mL; Thermo Fisher; PI62249) was used to mark nuclei. Live imaging was performed using an environmental control chamber warmed to 37°C and set to 5% CO_2_.

Prior to analysis, all images were denoised using Nikon AR analysis software (v.5.30.05). Representative images were shown upon automatic deconvolution using Nikon AR analysis software. Directional spread of stable microtubules was marked by alpha tubulin (acetyl K40) and quantified as relative frequency using the Directionality plugin in Fiji (ImageJ). Where indicated, mitochondria were observed upon transfection with eGFP-OMP25 plasmid for 24 h (125 ng/mL; Addgene # 14150), or live-cell staining with 100 nM MitoTracker^TM^ Red CMXRos (Thermo Fisher; M7512) for 15 min at 37°C prior to fixation or immediate imaging. Mitochondria count, volume, sphericity, and surface area were quantified in a minimum of 24 cells per condition using the Mitochondria Analyzer plugin in Fiji (ImageJ). Statistical significance was determined using the Kruskal-Wallis test in GraphPad Prism.

To tether the mitochondria and ER as previously described^49^, a plasmid bearing monomeric RFP fused to the outer mitochondrial membrane targeting sequence of mAKAP1 and the ER targeting sequence of yUBC6 (mAKAP1(34–63)-mRFP-yUBC6; gift from György Hajnóczky) was transfected for 24 h prior to imaging. Lipids were visualized upon staining cells with 2 µM BODIPY™ 493/503 or BODIPY™ 558/568 C_12_ (Thermo Fisher) as indicated for 1 h. Lipid droplet area was quantified in a minimum of 30 cells per condition using Fiji (ImageJ) and statistical significance was determined by two-way ANOVA and Tukey post hoc test in GraphPad Prism.

To quantify mitochondria-ER encapsulations (MENCs), cells were transfected with GFP-Sec61β (300 ng/mL; Addgene #15108) and stained with 100 nM MitoTracker^TM^ Red CMXRos (Thermo Fisher; M7512) for 15 min at 37°C. Images were taken every 5 sec for up to 5 min. MENCs were manually quantified as forming stable, asymmetric ER encapsulations (maintenance of contact ≥120 s; partial: maintenance of contact ≥45 s, < 120 s). Per condition, 20 mitochondria were quantified per cell in a minimum of 15 cells. Statistical significance was determined using a Welch’s t-test (\**P* < 0.05) in GraphPad Prism. To quantify mitochondrial calcium, cells were transfected with mitoCameleon (mtCameleon; pcDNA-4mtD3cpv; Addgene #36323). Z-slice images were taken live in a minimum of 10 cells per condition. Ratiometric cameleon signal was acquired via 405 nm excitation coupled with either 445 nm (CFP) or 488 nm (FRET) emission. Cameleon signal in all mitochondria per cell was quantified to determine the Rmin, Rmax, and % FRET ratio using the equation:

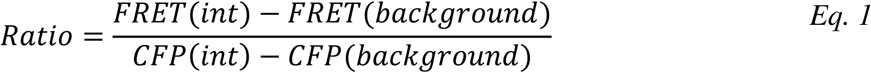

Statistical significance was determined using the Kruskal-Wallis test in GraphPad Prism.

### Oxygen Consumption Rate (OCR) and Extracellular Acidification Rate (ECAR)

OCR was determined using two methods. First, OCR and ECAR were measured using the Seahorse XF Cell Mito Stress Test Kit (Agilent) following the manufacturer’s protocol. Forty-eight hours prior to the Seahorse assay, a 37 °C incubator was set to 0% CO_2_. Twenty-four hours prior to the assay, the flux pack was hydrated with 200 µL of sterile H_2_O and placed within the 0% CO_2_ incubator. Four hours prior to the Seahorse assay, cells were plated into a pre-fibronectin-coated Agilent XFe96 plate. Two hours prior to the Seahorse assay, the water in the hydrated flux pack was replaced with prewarmed Seahorse XF Calibrant solution and returned to the 0% CO_2_ incubator for 1 h. After incubation, carbon sources and drugs were loaded into the plate. In port A was a mixed carbon source containing 10 mM glucose, 1 mM pyruvate and 2 mM glutamine. Ports B, C, and D contained 2 µM oligomycin, 1 µM FCCP and 0.5 µM Rotenone/Antimycin A, respectively. While calibrating this plate in the Seahorse analyzer, the cells were gently washed with Seahorse media and incubated at 0% CO_2_ for 1 h. OCR (pmol/min/cells) and ECAR

(mpH/min/cells) were continually analyzed following sequential carbon and drug treatment. Quantification of respiratory parameters non-mitochondria respiration, basal respiration, maximal respiration, proton leak ATP production and spare respiratory capacity were performed as previously described^36, 50^. Second, OCR was additionally monitored using the Resipher real-time 96-well plate oxygen sensor (Lucid Scientific). An equivalent number of cells per condition were seeded into a 96-well plate and the oxygen sensor was immediately placed onto the plate. OCR (fmol/mm^2^/sec) was monitored continuously for 16 h.

## Mass Spectrometry (MS) Analysis

### MS Sample Preparation-Whole Proteome and Parallel Reaction Monitoring (PRM) MS

Biological triplicates were processed for whole proteome analysis and targeted MS analysis by PRM. Organoids were collected from Matrigel upon incubation and three sequential washes with Cell Recovery Solution (Corning). Samples were pelleted, then lysed in 5% SDS and 50 mM triethylammonium bicarbonate, pH 8.5. Proteins were fully denatured in this solution upon bath sonication (20 bursts of 1 sec pulses at 60% output) and heating at 95°C for 5 min. BCA assay (Pierce) was used to determine protein concentrations. Ten micrograms protein per sample were reduced and alkylated with 25 mM tris(2-carboxyethyl)phosphine and 50 mM chloroacetamide upon incubation at 70°C for 20 min. Samples were acidified with aqueous phosphoric acid to a final concentration of 2.5% and loaded onto S-Trap micro spin columns (ProtiFi). Following the manufacturer’s protocol, S-Trap columns were used for protein binding, desalting, trypsin digestion, and peptide extraction. Trypsin digestion was performed at a 1:25 (w/w) enzyme:sample protein ratio and incubated overnight at 37°C. Peptides were eluted, dried down using a SpeedVac, and resuspended in UHPLC-MS water containing 1% formic acid, 4% UHPLC-MS acetonitrile, and 0.015% *n*-dodecyl β-D-maltoside.

#### Whole Proteome MS Data Acquisition

Peptides were analyzed by liquid chromatography and tandem mass spectrometry (LC-MS/MS) with a nanoElute 2 coupled to a timsTOF Ultra mass spectrometer (Bruker Daltonics). The mobile phases used were 0.1% formic acid in UHPLC-MS water (buffer A) and 0.1% formic acid/99.9% UHPLC-MS acetonitrile (buffer B). Peptides (100ng) were separated using a one column method, with the column heated to 50°C and attached to a CaptiveSpray Ultra ion source. A PepSep ULTRA C18 HPLC column (25 cm x 75 µm x 1.5 µm; Bruker) and a 10 µM emitter (Bruker) was used for samples in comparisons between 2D monolayers in 3D media and 3D organoids, as well as protein signatures of organoids, spheroids, and monolayers. Peptides from these samples were resolved using a linear gradient of 4% to 35% buffer B over 55 min at a flow rate of 200 nL/min. An IonOpticks Aurora Ultimate CSI C18 UHPLC packed emitter column (25 cm x 75 µm x 1.7 µm) was used for all other samples. Peptides from these samples were resolved using a linear gradient of 2% to 35% buffer B over 60 min at a flow rate of 200 nL/min.

For all samples, data-independent acquisition (DIA) mode with parallel accumulation serial fragmentation (dia-PASEF) was applied, containing 16 dia-PASEF scans separated into three ion mobility windows per scan (Supplementary Table 12). The MS scan range was 100 to 1700 m/z with positive ion polarity. TIMS settings included an ion mobility 1/K0 range of 0.65 to 1.46 V·s/cm^2^ and a 100 ms ramp time.

#### Whole Proteome MS Data Analysis

All DIA data were analyzed in DIA-NN^51, 52^ (version 1.9.2) with a mass spectral library containing a Uniprot mouse reference proteome (UP000000589 FASTA downloaded November 2024) and common contaminants. The maximum number of missed cleavages with trypsin was set to 1, N-terminal methionine excision enabled, fixed cysteine carbamidomethylation, precursor charge states of 1 to 4, peptide lengths of 7 to 30 amino acids, precursor ion *m/z* range from 300 to 1,800, and fragment ion *m/z* range from 200 to 1,800. The precursor false discovery rate (FDR) was set to 1.0%. The scan window radius was set to 7, and the MS1 and MS2 mass accuracies were set to 11 and 14 ppm, respectively. Cross-run normalization was disabled. DIA-NN results were analyzed using FragPipe-Analyst^53^ with minimum value imputation and Benjamini Hochberg FDR correction. Proteins with at least 2 unique peptides were quantified. For a given comparison of conditions, differentially abundant proteins were defined as having a log_2_ fold change value of |0.58| and an adjusted *p*-value of ≤0.05. Differentially abundant proteins were subjected to gene ontology, pathway enrichment, and cellular component analyses using STRING^54^ through the StringApp Plugin in Cytoscape^55, 56^ (version 3.10.3). Edges between protein nodes were determined based on high confidence and all protein interaction sources active except for textmining. k-means clustering was performed using Morpheus (Broad Institute; https://software.broadinstitute.org/morpheus). Tissue-specific functional module detection using HumanBase (Flatiron Institute – Simons Foundation)^57, 58^ was performed with a human oviduct background upon mouse to human gene name conversion. Protein constituents of Matrigel® were determined as previously described^59^.

#### PRM MS Data Acquisition

Proteotypic peptides for MCS proteins were prioritized as follows. First, proteotypic peptides identified in a whole proteome MS analysis using data-dependent acquisition (DDA) mode were retained. Second, mouse orthologs of human proteotypic peptides for each MCS protein, which were previously identified by PRM^30^, were retained.

DDA and PRM LC-MS/MS analysis of peptides (1.5 µg) was performed on a Dionex Ultimate 3000 nanoRSLC coupled online to an EASYSpray ion source and a Q Exactive HF Hybrid Quadrupole-Orbitrap mass spectrometer (Thermo Fisher Scientific). Peptides were separated on an EASYSpray PepMap RSLC C18 column (75 µm x 25 cm) heated to 50°C. The mobile phases used were 0.1% formic acid in UHPLC-MS water (buffer A) and 0.1% formic acid/99.9% UHPLC-MS acetonitrile (buffer B). The linear gradients performed in the DDA run were 5% to 22% buffer B over 33 min, followed by 22% to 40% over 27 min at a 250 nL/min flow rate. The linear gradients performed in the PRM were 2% to 22% buffer B over 45 min, followed by 22% to 38% over 15 min at a 250 nL/min flow rate. MS1 scans were acquired at a resolution of 120,000, automatic gain control (AGC) target of 3e6, maximum inject time (MIT) of 30 ms, and a 350 to 1800 *m/z* scan range. The top 20 precursors were selected for fragmentation. MS2 scans were acquired at a resolution of 15,000, AGC target of 1e5, MIT of 25 ms, isolation window of 1.2 *m/z*, fixed first mass of 100.0 *m/z*, normalized collision energy of 28, and dynamic exclusion of 45 sec.

#### PRM MS Data Analysis

RAW files were imported into Skyline^60, 61^ to extract the chromatograms and calculate peak areas of the product ions. Proteome Discoverer was also used to search the RAW files, and the search results were imported as a spectral library into Skyline. Spectra were searched against a Uniprot mouse reference proteome (UP000000589) appended with common contaminants. The SEQUEST HT algorithm was used to search for tryptic peptides with ≤ 2 missed cleavages, a precursor mass tolerance of 10 ppm, and a fragment mass tolerance of 0.02 Da. Percolator was used to score the matched spectra. In Skyline, peptides were prioritized to maintain the ≥ 3 highest intensity fragment ions with the lowest mass errors. Peak areas for each peptide were summed and normalized to housekeeping H2B1B and H2A2C peptides. Peptide abundance comparisons were calculated as the indicated ratios, and proteins with at least 2 unique peptides were averaged per replicate, and then averaged across replicates. Individual values of each protein per sample were plotted and statistical significance was determined with a Mann-Whitney test in GraphPad Prism.

## Acknowledgments.

This study was supported by funding from Stand Up To Cancer Convergence 3.1416 (to I.M.C. and

B.G.N.), NIH (R01AI174515 to I.M.C. and R01CA257507 to B.G.N.), Paul G. Allen Family Foundation (to I.M.C.), Pershing Square Foundation Ovarian Cancer grant (to B.G.N.), a NCATS TL1 award (TL1TR003019 to K.K.L), and an American Heart Association fellowship (23PRE1014367 to W.A.H.).

## Author Contributions

Conceptualization, K.K.L. and I.M.C.; investigation, K.K.L., O.J.P., W.A.H., J.M.M., and I.A.J.; formal analysis, K.K.L.; methodology, K.K.L., O.J.P., and B.G.N.; data curation, K.K.L.; visualization, K.K.L.; resources, K.K.L., O.J.P., I.M.C., and B.G.N.; validation, K.K.L.; O.J.P., and W.A.H.; writing – original draft, K.K.L. and I.M.C.; writing – review & editing, K.K.L., I.M.C., O.J.P., and B.G.N.; funding acquisition, I.M.C. and B.G.N.; supervision, I.M.C. and B.G.N.; project administration, I.M.C. and B.G.N.

## Declaration of generative AI and AI-assisted technologies in the writing process

The authors declare that no generative AI and AI-assisted technologies were used in the writing process.

## Supplementary Table Legends

**Supplementary Table 1.** Protein abundances of *Trp53^-/-^*;*Pten^-/-^*;*Nf1^-/-^*(PPN), and *Trp53^-/-^;Brca1^+/-^;Myc^OE^* (PBM) models grown as 3D organoids in 3D media.

**Supplementary Table 2.** Functional modules enriched in HGSC states relative to parental organoids (PPN versus P and PBM versus P). Functional enrichment was determined using HumanBase with mouse to human gene ortholog conversion and a tissue-specific oviduct background.

**Supplementary Table 3.** k-means clustering and GO enrichment of proteins across PPN and PBM in 2D and 3D conditions.

**Supplementary Table 4.** Proteomic analysis of functional modules underrepresented in PPN relative to PBM organoids as determined by HumanBase with mouse to human gene ortholog conversion.

**Supplementary Table 5.** Protein abundance comparison of HGSC models grown as 2D monolayers, 3D organoids, and 3D spheroids. Samples were grown in 2D and 3D media as indicated.

**Supplementary Table 6.** Proteomic analysis of GO Biological process terms underrepresented in 3D organoids relative to 2D monolayers grown in 3D media.

**Supplementary Table 7.** Proteomic analysis of GO Cellular component terms underrepresented in 3D organoids relative to 2D monolayers grown in 3D media.

**Supplementary Table 8.** Proteomic analysis of GO Biological process terms overrepresented in 3D organoids relative to 2D monolayers grown in 3D media.

**Supplementary Table 9.** Proteomic analysis of GO Cellular component terms overrepresented in 3D organoids relative to 2D monolayers grown in 3D media.

**Supplementary Table 10.** Cellular component enrichment of lipid-associated proteins elevated in HGSC *Trp53^-/-^*;*Pten^-/-^*;*Nf1^-/-^* (PPN) relative to *Trp53^-/-^;Brca1^+/-^;Myc^OE^* (PBM) (derived from Cluster 1 in Figure 1C-D).

**Supplementary Table 11.** Membrane contact site (MCS) protein and accompanying peptide abundances quantified by parallel reaction monitoring in HGSC subtypes grown as 2D monolayers and 3D organoids. MCS proteins were categorized as mediating functional contacts between mitochondria-endoplasmic reticulum (ER), mitochondria-lysosome (lyso), ER-endolysosome (endolyso), ER-Golgi, ER-plasma membrane (PM), peroxisome (PEX)-ER, and outer mitochondrial membrane – inner mitochondrial membrane (mito OMM-IMM).

**Supplementary Table 12**. Variable isolation windows used in the mass spectrometry dia-PASEF method.

