## Supplementary Figures for "Monolayer, Spheroid, and Organoid Formats Elicit Proteomic States that Couple Organelle Organization to Metabolic Vulnerabilities"

**Supplementary Figure 1**.

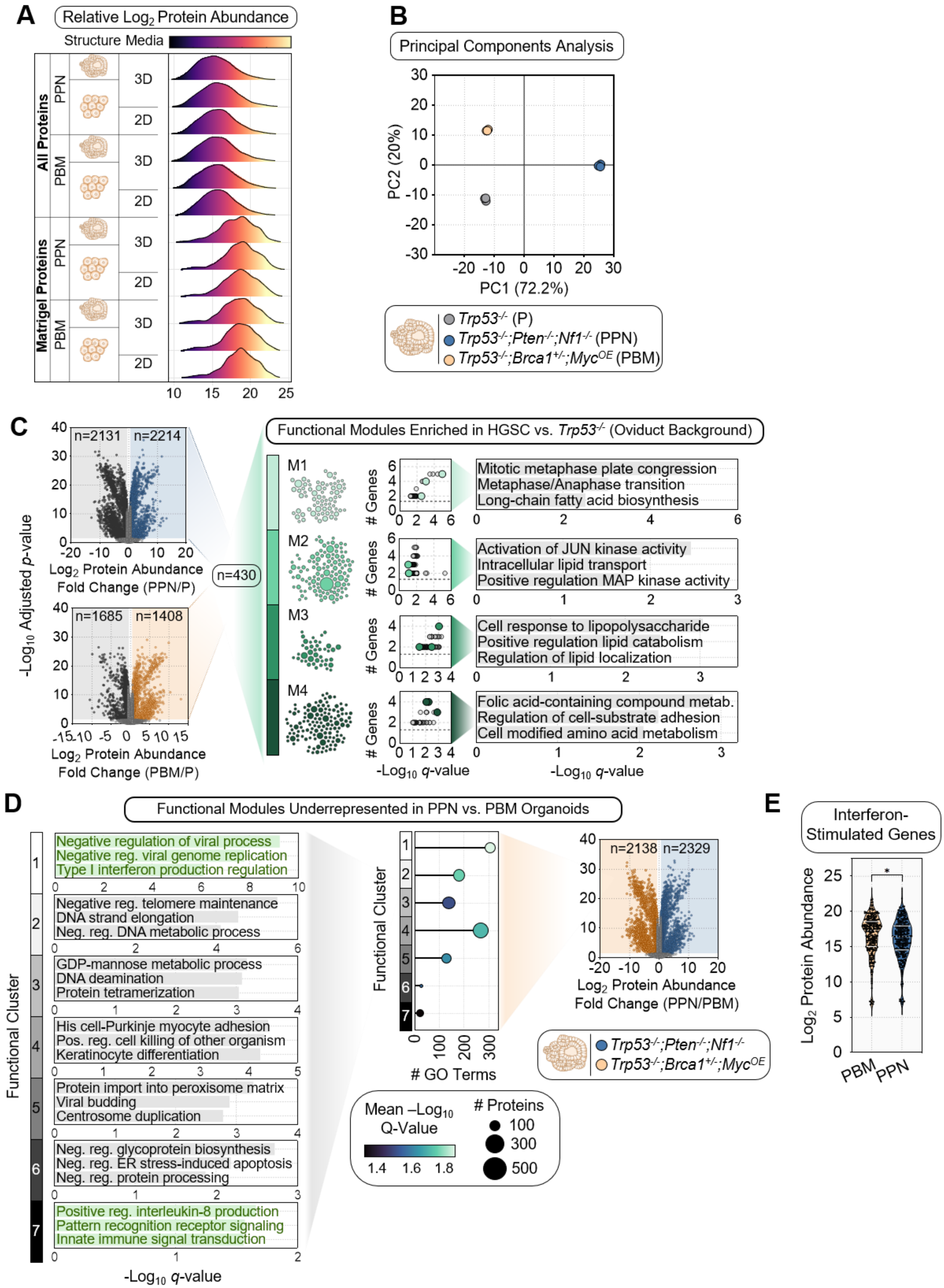

**Supplementary Figure 1. HGSC cells diverge from the parental cells and differ in basal antiviral programs.**

(A) Relative abundance distributions for all quantified proteins, and proteins reported to comprise Matrigel® in HGSC genotypes *Trp53*$^{-/-}$;*Pten*$^{-/-}$;*Nf1*$^{-/-}$ (PPN), and *Trp53*$^{-/-}$;*Brca1*$^{+/-}$;*Myc*$^{OE}$ (PBM), and growth formats.

(B) Principal components analysis of parental *Trp53*$^{-/-}$ (P), PPN, and PBM organoid proteomes.

(C) Genotype-resolved proteomic comparisons of PPN versus P and PBM versus P cells, showing volcano plots and representative functional modules enriched in HGSC cells relative to parental organoids. Functional enrichment was determined using HumanBase[37] with mouse to human gene ortholog conversion and a tissue-specific oviduct background.

(D) Functional modules underrepresented in PPN relative to PBM organoid proteomes as determined by HumanBase, including a volcano plot, representative pathways, and differential abundance summary.

(E) Basal abundance of proteins encoded by interferon-stimulated genes in PBM and PPN organoids.

**Supplementary Figure 2.**

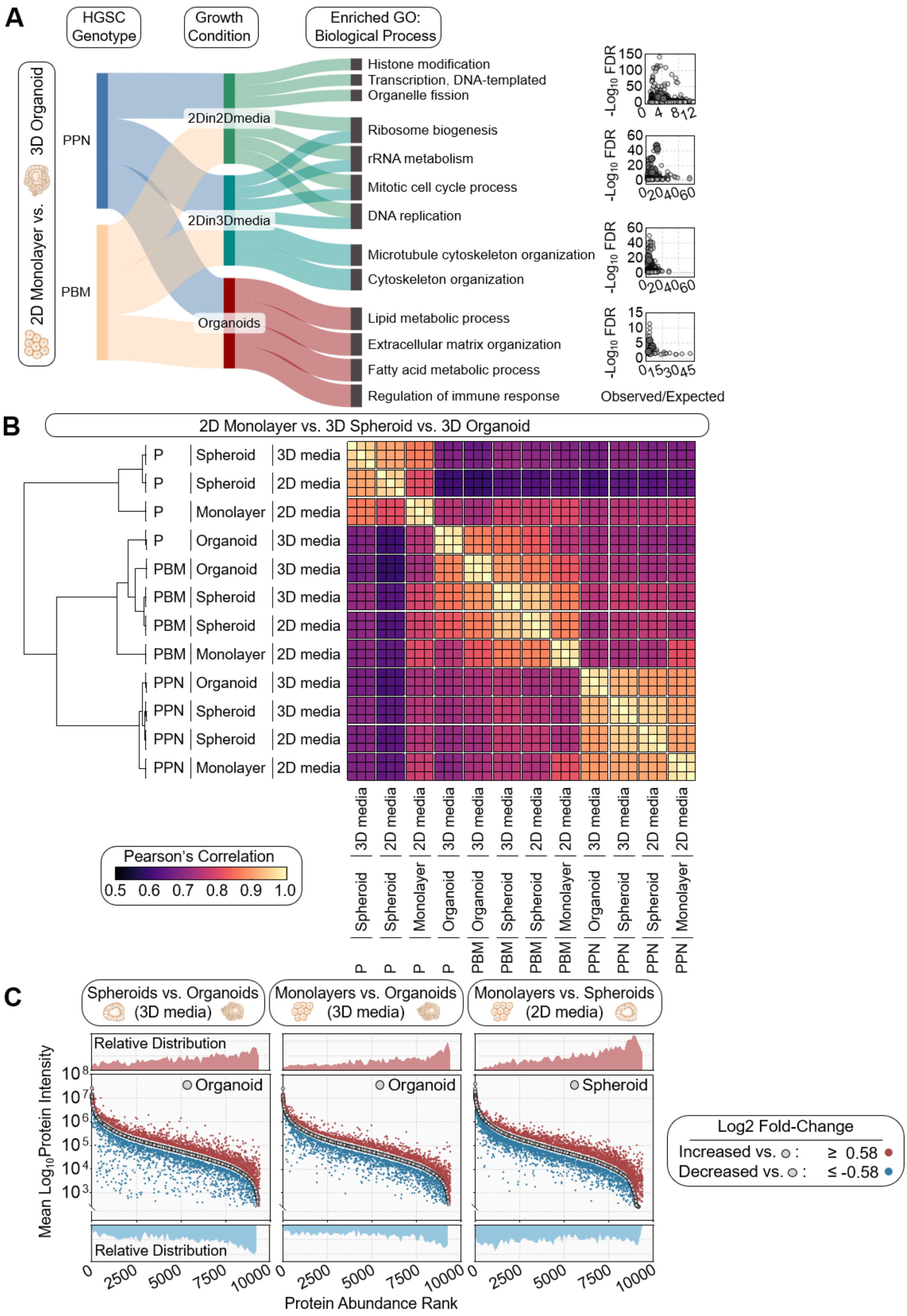

**Supplementary Figure 2. Analyses supporting distinct architecture- and media-dependent proteomic states.**

(A) Process-level summary of enriched GO biological processes distinguishing 2D monolayers versus 3D organoids across *Trp53*$^{-/-}$;*Pten*$^{-/-}$;*Nf1*$^{-/-}$ (PPN) and *Trp53*$^{-/-}$;*Brca1*$^{+/-}$;*Myc*$^{OE}$ (PBM) models.

(B) Pearson correlation and hierarchical clustering of parental *Trp53*$^{-/-}$ (P), PPN, and PBM monolayer, spheroid, and organoid cultures in 2D or 3D media, as indicated.

(C) Rank-ordered protein abundance plots comparing spheroids and organoids (3D media), monolayers and organoids (3D media), and monolayers and spheroids (2D media). Proteins increased (red; $\log_2$ fold-change ≥ 0.58) or decreased (blue; $\log_2$ fold-change ≤ -0.58) are highlighted.

**Supplementary Figure 3.**

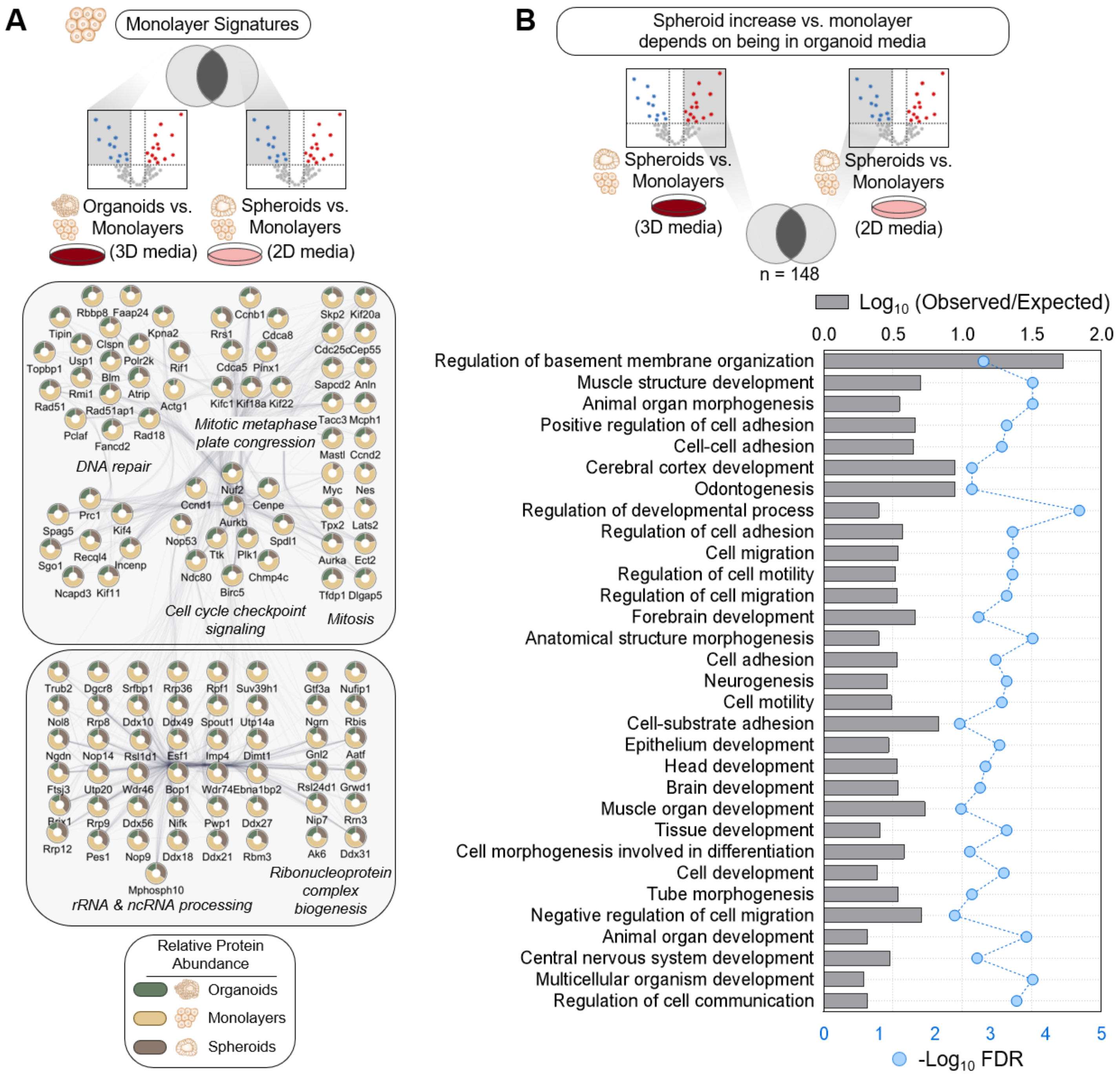


**Supplementary Figure 3. Monolayer-enriched and media-conditioned spheroid-associated proteome analyses.**

(A) Protein-protein interaction networks of monolayer-enriched protein signatures. Proteins were annotated using the STRING database. Nodes display relative abundance in monolayers, spheroids, and organoids. Edges represent functional protein interactions.

(B) Analysis of proteins whose spheroid-versus-monolayer increase depends on organoid media, with representative enriched GO biological process terms annotated using the STRING database.

**Supplementary Figure 4.**

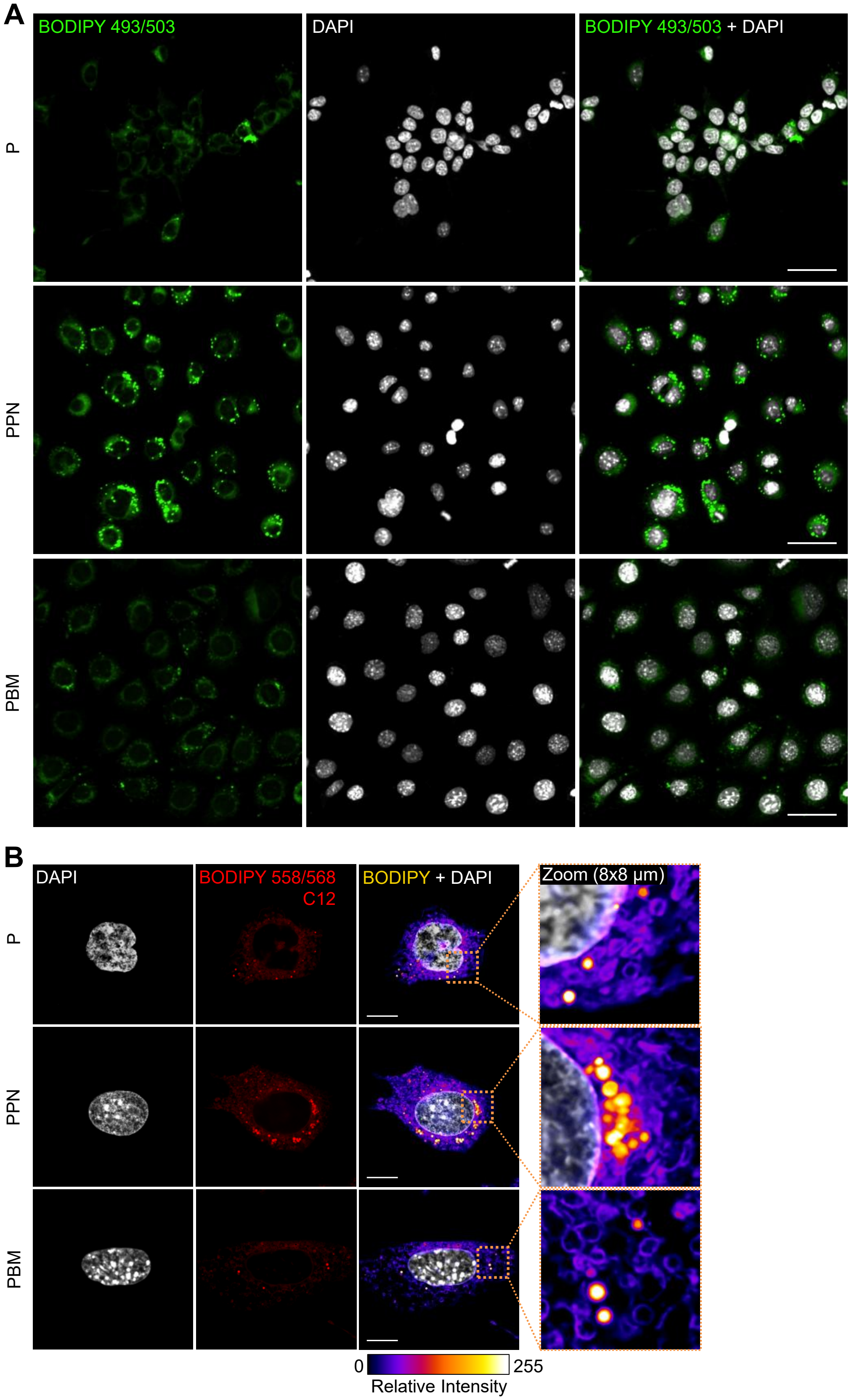


**Supplementary Figure 4. Subcellular lipid organization in parental and HGSC cells.**
(A) Representative BODIPY 493/503 staining of parental *Trp53*$^{-/-}$ (P), *Trp53*$^{-/-}$;*Pten*$^{-/-}$;*Nf1*$^{-/-}$ (PPN), and *Trp53*$^{-/-}$;*Brca1*$^{+/-}$;*Myc*$^{OE}$ (PBM) cells. Scale bar: 50 µm.
(B) Representative BODIPY 558/568 C12 staining across parental P, PPN, and PBM cells, with merged views and zoomed regions highlighting the organization of lipid droplet clusters. Scale bar: 10 µm.

**Supplementary Figure 5.**

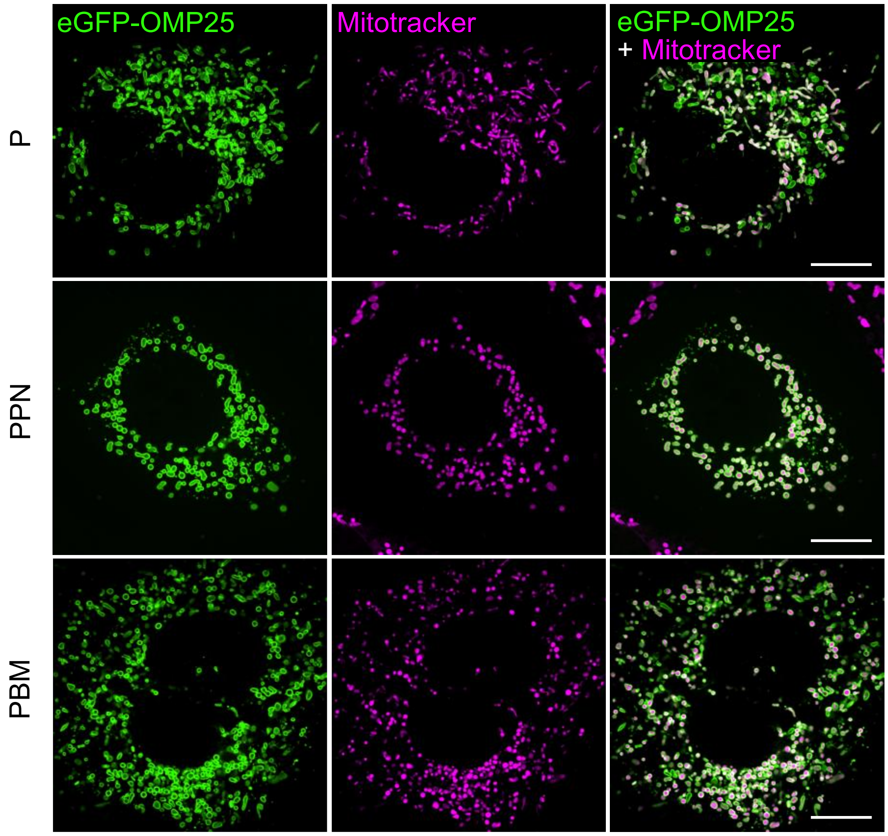


**Supplementary Figure 5. HGSC genotype-dependent mitochondrial remodeling.**
Representative confocal imaging of eGFP-OMP25 and MitoTracker Red CMXRos views in parental *Trp53*$^{-/-}$ (P), *Trp53*$^{-/-}$*;Pten*$^{-/-}$*;Nf1*$^{-/-}$ (PPN), and *Trp53*$^{-/-}$*;Brca1*$^{+/-}$*;Myc*$^{OE}$ (PBM) cells, showing mitochondrial labeling and subtype-dependent differences in mitochondrial organization.

**Supplementary Figure 6.**

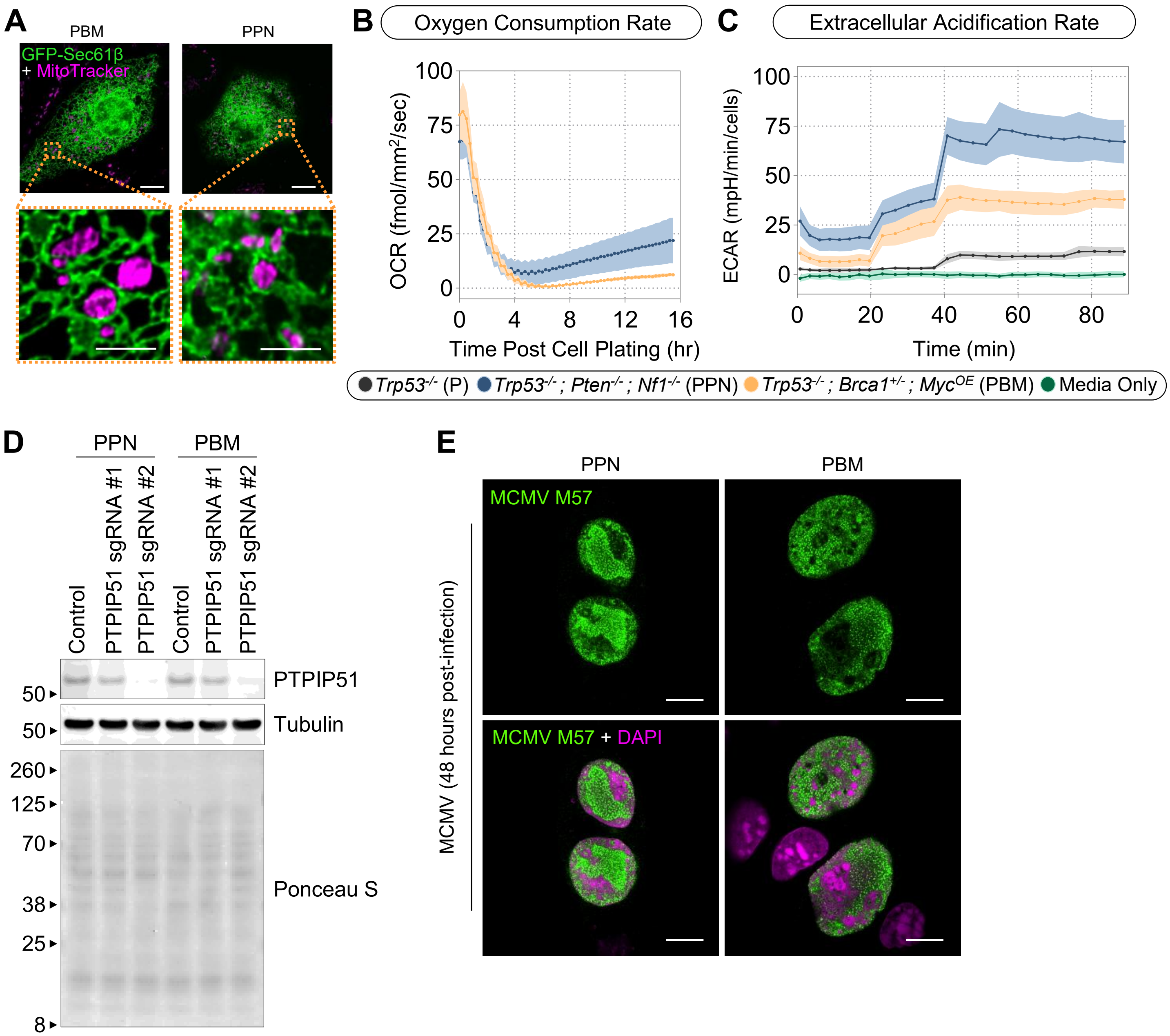


**Supplementary Figure 6. Functional implications of increased mitochondria-ER contacts in PPN cells.**
(A) Representative confocal imaging of ER (GFP-Sec61β) and mitochondria (MitoTracker Red CMXRos) in *Trp53*$^{-/-}$;*Pten*$^{-/-}$;*Nf1*$^{-/-}$ (PPN) and *Trp53*$^{-/-}$;*Brca1*$^{+/-}$;*Myc*$^{OE}$ (PBM) cells, with zoomed-in regions highlighting differences in mitochondria-ER contacts.
(B) Oxygen consumption rates (OCR) in parental *Trp53*$^{-/-}$ (P), PPN, and PBM cells cultured in 2D media. A Resipher oxygen sensor was used.
(C) Seahorse analysis showing extracellular acidification rates (ECAR) in parental P, PPN, and PBM cells.
(D) Immunoblot validation of PTPIP51 depletion in PBM and PPN cells using independent sgRNAs and CRISPR/Cas9 gene editing.
(E) Representative confocal imaging of MCMV M57 staining at 48 h post-infection in PPN and PBM cells. MCMV M57 is the viral major single-stranded DNA binding protein.
